# Remyelination protects neurons from DLK-mediated neurodegeneration

**DOI:** 10.1101/2023.09.30.560267

**Authors:** Greg J. Duncan, Sam D Ingram, Katie Emberley, Jo Hill, Christian Cordano, Ahmed Abdelhak, Michael McCane, Jennifer E. Jenks, Nora Jabassini, Kirtana Ananth, Skylar J. Ferrara, Brittany Stedelin, Benjamin Sivyer, Sue A. Aicher, Thomas Scanlan, Trent A. Watkins, Anusha Mishra, Jonathan W. Nelson, Ari J. Green, Ben Emery

## Abstract

Chronic demyelination and oligodendrocyte loss deprive neurons of crucial support. It is the degeneration of neurons and their connections that drives progressive disability in demyelinating disease. However, whether chronic demyelination triggers neurodegeneration and how it may do so remain unclear. We characterize two genetic mouse models of inducible demyelination, one distinguished by effective remyelination and the other by remyelination failure and chronic demyelination. While both demyelinating lines feature axonal damage, mice with blocked remyelination have elevated neuronal apoptosis and altered microglial inflammation, whereas mice with efficient remyelination do not feature neuronal apoptosis and have improved functional recovery. Remyelination incapable mice show increased activation of kinases downstream of dual leucine zipper kinase (DLK) and phosphorylation of c-Jun in neuronal nuclei. Pharmacological inhibition or genetic disruption of DLK block c-Jun phosphorylation and the apoptosis of demyelinated neurons. Together, we demonstrate that remyelination is associated with neuroprotection and identify DLK inhibition as protective strategy for chronically demyelinated neurons.

**Highlights:**

- Characterization of a transgenic mouse model of demyelination without subsequent remyelination
- Remyelination protects neurons from axon loss and neuronal apoptosis
- MAPK and c-Jun phosphorylation are increased in mice featuring remyelination failure
- DLK is necessary for the apoptosis of chronically demyelinated neurons

## Introduction

Remyelination is the regenerative process by which new myelin sheaths are produced in the CNS, typically via the differentiation of oligodendrocytes (OLs) from oligodendrocyte precursor cells (OPCs)^1, 2^, or to a limited extent by OLs that survive the demyelinating insult^3, 4, 5^. In the inflammatory demyelinating disease multiple sclerosis (MS), remyelination is often incomplete^6, 7^, resulting in chronic demyelination of axons. Chronic demyelination and the loss of OLs deprive neurons of crucial metabolic and trophic support and is hypothesized to leave them vulnerable to subsequent degeneration^8, 9, 10^. With disease chronicity, a progressive loss of neurons and their axons in MS drives permanent disability^11, 12^. While several studies have found an association between poor remyelination efficiency and neurodegeneration in MS^13, 14^, there is a paucity of experimental evidence demonstrating that impaired oligodendrogenesis and remyelination failure lead to neurodegeneration. To date, no study has selectively targeted the OL lineage to induce demyelination and determined the extent of neurodegeneration in the context of subsequent successful or failed remyelination. Additionally, the molecular mechanisms critical for neurodegeneration following demyelination remain unclear

Here, we compare two rodent models of genetic demyelination, which feature either successful or impaired remyelination. By contrasting these two rodent models, we find that mice with successful remyelination do not feature axonal degeneration and neuronal apoptosis in the visual system, whereas mice unable to remyelinate have fewer axons and increased retinal ganglion cell (RGC) apoptosis. Mice with failed remyelination have elevated phosphorylation of kinases downstream of DLK in the optic nerve and phosphorylation of the transcription factor c-Jun in RGC nuclei, suggesting that DLK activation promotes degeneration of demyelinated neurons. Pharmacological inhibition or genetic disruption of DLK block both the phosphorylation of c-Jun and RGC apoptosis. We propose that effective remyelination of the axon improves neuronal survival by suppressing DLK-mediated signaling.

## Results

### Myelin regulatory factor (*Myrf*) knockout from both OPCs and mature OLs results in genetic demyelination and impaired remyelination

A rodent model that inhibits remyelination in a cell-selective manner would permit an understanding of the extent and mechanisms by which prolonged demyelination damages neurons. Myrf is an essential transcription factor for myelin gene transcription^15, 16^, and its deletion from mature OLs using Plp1-CreERT mice (Myrf^fl/fl^ Plp1 CreERT; hereto referred to as Myrf^ΔiPlp1^) results in CNS-wide demyelination^17^. Reasoning that extending the knockout of *Myrf* to OPCs would prevent remyelination^18^, we crossed the Myrf^fl/fl^ line to the pan-OL lineage Sox10-CreERT^19^ line (Myrf^fl/fl^ Sox10 CreERT; Myrf^iSox10^) (Fig. 1a). Dosing of both Myrf^ΔiPlp1^ and Myrf^ΔiSox10^ mice with tamoxifen at eight weeks of age (Fig. 1b) results in progressive ataxia, hindlimb tremor, and paresis (Supplementary Fig.1a,b) coincident with CNS-wide demyelination (Supplementary Fig.1c,d). In Myrf^ΔiPlp1^ mice these symptoms peaked by 10-12 weeks post-treatment and gradually improved, as previously published^17, 20^. In contrast, Myrf^ΔiSox10^ mice had reduced mobility, weight loss and after 12 weeks post tamoxifen developed seizures, necessitating their euthanasia (Supplementary Fig.1a,b).

**Fig. 1.**
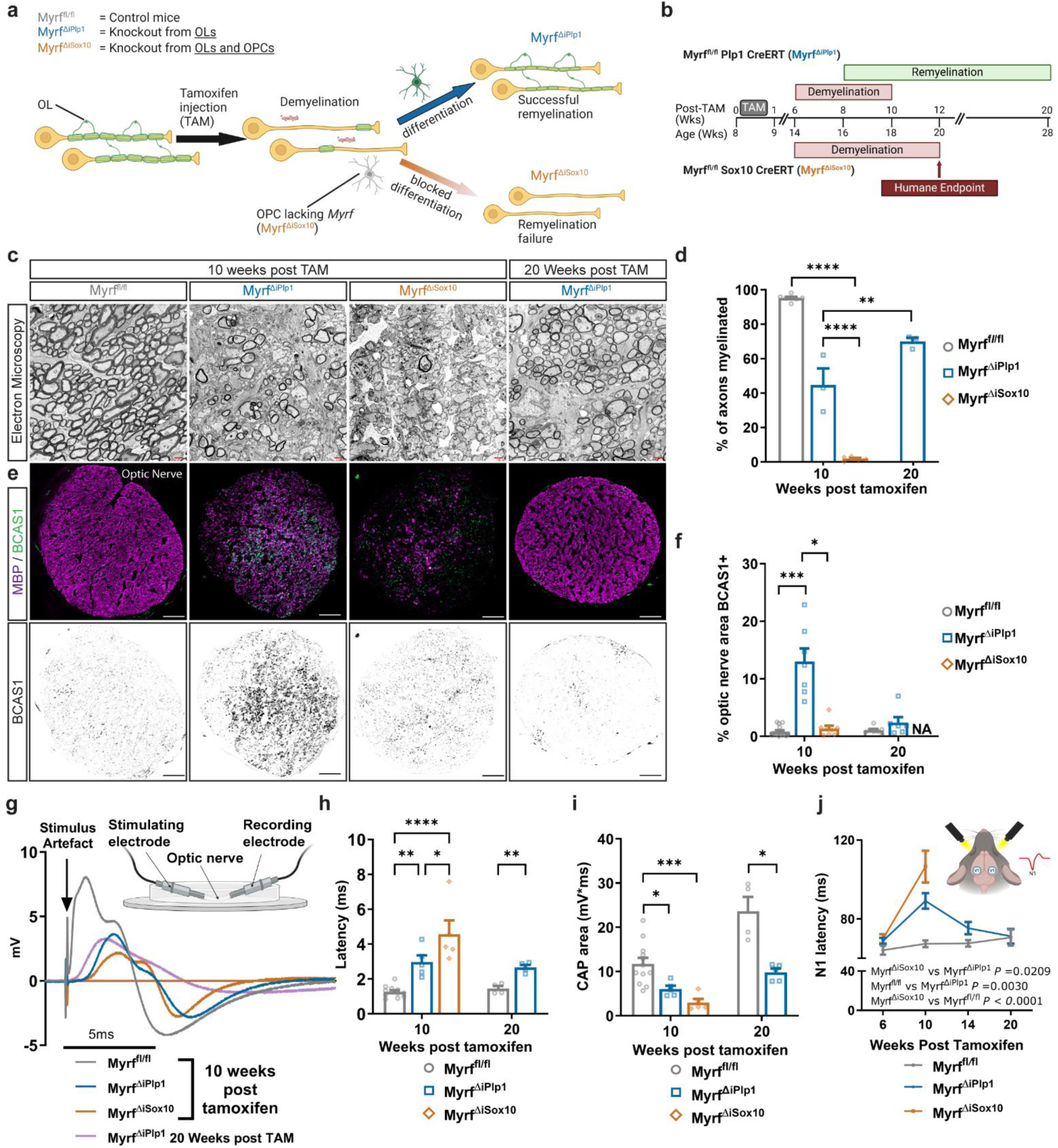
Myrf^ΔiSox10^ mice undergo CNS demyelination with limited remyelination. **a** Transgenic strategy to induce demyelination with the aim of permitting (Myrf^ΔlPlp1^ mice) or inhibiting (Myrf^ΔiSox10^ mice) remyelination, b Timeline of tamoxifen administration, demyelination and remyelination in Myrf^Δiplp1^ and Myrf^ΔiSox10^ mice, **c** Electron micrographs of the optic nerve of Myrf^fl/fl^, Myrf**^ΔiPIp1^** and Myrf^ΔiSox10^ mice, **d** Quantification of the percentage of myelinated axons in the optic nerve. There is a decline in the percentage of axons myelinated in ^ΔiSox10^ and Myrf^aiplp1^relative to Myrf^fl/fl^mice at 10 weeks post tamoxifen (*P* = 0.0001). MyrP^ΔiSox10^ mice have increased myelinated axons week at 20 weeks relative to 10 weeks post tamoxifen *(P* = 0.0017). e Optic nerve cross sections stained for MBP (myelin) and BCAS1 (new OLs/myelin). **f** Percentage of optic nerve that is BCAS1+. At 10 weeks post tamoxifen, Myrf^,ΔiPlp1^ mice have increased BCAS1 relative to Myrf^fl/fl^ mice *(P-* 0.0007) and Myrf^ΔiSox10^ mice (*P* = 0.0109). **g** Example CAPs from the optic nerve, **h** CAP latency during de- and remyelination. Both Myrf^ΔiPlp1^ (*P* = 0.0069) and Myrf^ΔiSox10^(*P* = 0.0001) nerves have increased latency at 10 weeks post-tamoxifen relative to non-demyelinated Myrf^fl/fl^ nerves. Myrf^ΔiSox10^ nerves have increased latency relative to Myrf^ΔiPlp1^ nerves at week 10 post tamoxifen *(P=* 0.0342). By 20 weeks post tamoxifen, Myrf^ΔiPlp1^ nerves do not fully recover in latency relative to Myrt^fl/fl^ *(P-* 0.0012). **i** CAP area during de- and remyelination. Both Myrf^ΔiPlp1^ (*P* = 0.0287) and Myrf^ΔiSox10^ (*P* = 0.0010) nerves have decreased CAP area relative to Myrf^fl/fl^ nerves at 10 weeks post tamoxifen. By 20 weeks post tamoxifen, Myrf^ΔiPlp1^ nerves still have reduced CAP area relative to Myrf^fl/fl^ *(P=* 0.0205). **j** VEPs measured over time Myrf^ΔiPlp1^ (*P* = 0.0030) and Myrf^ΔiSox10^ (P< 0.0001) differed from Myrf^fl/fl^ and Myrf^ΔiPlp1^ had decreased latency relative to Myrf^ΔiSox10^ (*P* = 0.0209). One-way ANOVA with Tukey’s *post hoc* for **d**, and used for week 10 h and I. Kruskal Wallis with Dunn’s test for **d** and repeated measures mixed effect model with Tukey’s post hoc for **j.** Week 20 comparisons used Student’s t-test for **h** and with Welch correction for **i.** Mann Whitney U test was used for week 20 comparison in **e.** Scale bar is 1pm in **c,** 50 pm in **e.** NA = Not applicable. Error bars are SEM.

We took advantage of the optic nerve, a structure comprised of RGC axons that are nearly all myelinated to compare the remyelination potential of each mouse line (Fig. 1c). By 10 weeks post tamoxifen, only 44.7 ± 9.5% of axons in the optic nerves of Myrf^ΔiPlp1^ mice were myelinated, with the majority of the myelin sheaths present being thin (g-ratio > 0.80), characteristic of remyelination (Supplementary Fig.1e-h). By 20 weeks post tamoxifen, the degree of myelination increased to 75% of the axons in the optic nerve of Myrf^ΔiPlp1^ mice (Fig. 1d). In contrast, only 1.8 ± 0.6% of axons Myrf^ΔiSox10^ mice were myelinated at 10 weeks post tamoxifen with almost no signs of thinly myelinated axons (Fig. 1c,d). Consistent with a failure of remyelination in the Myrf^ΔiSox10^ line, Myrf^ΔiPlp1^ but not Myrf^ΔiSox10^ mice showed elevated expression of BCAS1, a marker of newly-formed OLs and their myelin^21^, at 10 weeks post tamoxifen (Fig. 1e,f).

To assess the functional consequences of remyelination failure, we performed compound action potential (CAP) recordings of the optic nerve (Fig. 1g). There is an increased CAP onset latency and reduced CAP area in both demyelinated mouse lines (Fig. 1h,i). However, Myrf^ΔiPlp1^ mice show improved conduction speeds relative to Myrf^ΔiSox10^ mice at 10 weeks post tamoxifen (Fig. 1g,h). Similarly, visual evoked potentials, sensitive to remyelination efficacy^22^, are slower in Myrf^ΔiSox10^ mice relative to Myrf^ΔiPlp1^ mice, and Myrf^ΔiPlp1^ mice return to non-demyelinated control levels by 20 weeks post tamoxifen (Fig. 1j). In summary, Myrf^ΔiPlp1^ mice feature rapid and effective remyelination, which is associated with greater functional and electrophysiological recovery, whereas Myrf^ΔiSox10^ mice have little remyelination and worsened functional outcomes.

### New oligodendrocytes are unable to fully mature in Myrf^ΔiSox10^ mice

To better understand the cellular changes seen in each model of demyelination, we next performed single-nuclei RNA sequencing (snRNA-seq) of the optic nerves of Myrf^fl/fl^, Myrf^ΔiPlp1^, and Myrf^ΔiSox10^ mice (Fig. 2a). 49,806 nuclei passed quality control and were sorted into 14 distinct clusters based on well-characterized markers (Fig. 2b-d). Both Myrf^ΔiPlp1^ and Myrf^ΔiSox10^ mice show a near complete loss of the mature OL (MOL) cluster (Fig. 2e,f), consistent with efficient recombination and near absence of the developmentally-generated OLs in both lines. Myrf^ΔiPlp1^ mice had increased proportions of both the newly-formed OL (NFOL, characterized by high *Enpp6, Bcas1 and Tcf7l2*) and myelin-forming OL (MFOL, characterized by high expression of *Mobp, Mbp* and other myelin protein transcripts) populations. Together, this supports active oligodendrogenesis and remyelination in Myrf^ΔiPlp1^ mice (Fig. 2f). In contrast, Myrf^ΔiSox10^ mice formed virtually no NFOLs or MFOLs and lacked expression of myelin protein transcripts (*Mobp, Mog, Mag*) and critical genes for OL function like *Anln* and *Trf* (Fig. 2g). The Myrf^ΔiSox10^ mice form committed oligodendrocyte precursors (COPs), which are an intermediate differentiation state characterized by the downregulation of OPC markers *Pdgfrα* and *Cspg4* and the expression of *Nkx2.2* and *Fyn (*Supplementary Data Fig. 2c). However, these cells in the Myrf^ΔiSox10^ mice lack the characteristic expression of *Gpr17* and cluster separately (COP2) from the COPs seen in Myrf^ΔiPlp1^ mice (COP1) (Supplementary Fig. 2a-c). COP1 cells are transcriptionally similar to NFOLs whereas COP2 share many transcripts with the knockout OL (KOOL) cluster (Supplementary Fig. 2c).

**Fig. 2.**
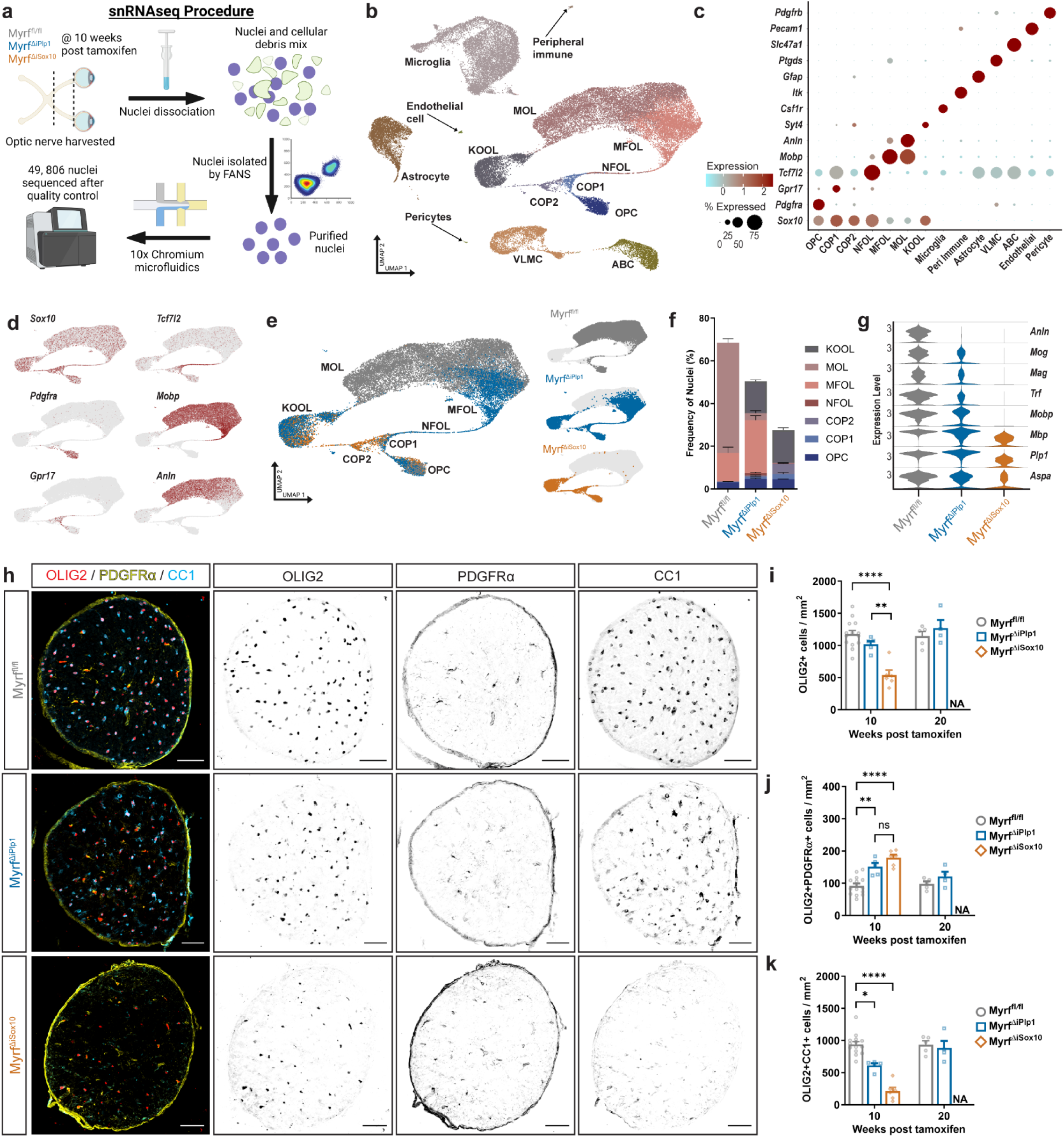
Differentiation of OPCs is blocked at the COP stage following demyelination in Myria^ΔiSox10^ mice. **a** Schematic of approach used to isolate and sequence nuclei from the optic nerve, **b** Uniform manifold approximation and projection (UMAP) of 49,806 nuclei. COP = committed OL precursor cells, NFOL = newly-formed OL, MFOL = myelin-forming OLs, MOL = mature OLs, KOOL = knockout OLs, VLMC = vascular and leptomeningeal cells ABC = arachnoid barrier cells, **c** Dot plot showing cluster-specific markers, **d** UMAP displaying expression of key markers of the OL lineage *(SoxIO),* OPCs *(Pdgfra),* COP1s *(Gpr17),* NFOLs *(Tcf7l2),* MFOLs *(Mobp)* and MOLs *(Anin, Mobp)* **e** UMAP of OL lineage cell nuclei broken down by genotype, **f** Stacked bar graph showing the proportions of oligodendroglial cell clusters across genotypes, **g** Violin plots for selected transcripts expressed in OL lineage cells across genotypes, h Optic nerve cross sections stained with the OL lineage marker OLIG2, along with PDGFRa (expressed in OPCs) and CC1 (in OLs). **i** The density of OLIG2+ oligodendrocyte lineage cells declines in Myrf^ΔiSox10^ mice at 10 weeks post tamoxifen relative to Myrf^ΔiPlp1,^ (*P* = 0.0013) and Myrf^fl/fl^ *(P* = 0.0001). j The density of OLIG2+PDGFRa+ OPCs increases in both Myrf^ΔiPlp1^ (*P* = 0.0016) and Myrf^ΔiSox10^ *(P<* 0.0001) mice at 10 weeks post-tamoxifen relative to Myrf^fl/fl^ control mice, but do not differ from each other (P= 0.2529). kThe density of OLIG2+CC1 + OLs decreases in both in both Myrf^ΔiPlp1^ *(P* = 0.0397) and Myrf^ΔiSox10^*(P* < 0.0001) at 10 weeks post-tamoxifen relative to Myri^fl/fl^ mice One-way ANOVA with Tukey’s host hoc for week 10 pairwise comparisons in i and j with Kruskal Wallis with Dunn’s test in **k.** Week 20 comparisons used Student’s t-test in **i**, and **j** and Mann Whitney U in **k**. Error bars are SEM. Scale bar is 50 pm in **h**. NA = Not applicable, ns = not statistically significant.

One unexpected observation is the formation of a distinct cluster of cells in the two *Myrf* conditional knockout lines, referred to as KOOLs. KOOLs express OL-enriched transcription factors like *Zfp536* and *St18* but also OPC/COP transcription factors like *Sox6*, *Zeb1,* and *Klf6* (Supplementary Fig. 3a). KOOLs also fail to robustly express key myelin genes or OPCs markers like *Cspg4* and *Pdgfra* (Supplementary Fig. 3a,d). KOOLs are produced in similar proportions in both Myrf^ΔiPlp1^ and Myrf^ΔiSox10^ when examined as the percentage of nuclei by snRNAseq (Supplementary Fig. 3f) or using SYT4 immunohistochemistry (Supplementary Fig. 3e,g and h). The KOOL population shares some markers with demyelination/disease-associated OLs present in cuprizone and Alzheimer’s disease, including *Col5a3, Serpina3n, Klk8* and *Cdkn1a*^23, 24, 25^ transcripts indicative of damage (Supplementary Fig. 3a,d). The presence of KOOLs at 10 weeks post tamoxifen suggests that OL lineage cells may persist for some time following *Myrf* ablation before undergoing apoptosis.

To validate the snRNAseq results, we performed immunohistochemistry on optic nerves (Fig. 2h). Total OL lineage cells (OLIG2+) declined selectively in Myrf^ΔiSox10^ mice (Fig. 2i) in large part due to the reduction of mature OLs (OLIG2+CC1+, Fig. 2k). In Myrf^ΔiPlp1^ mice, OL density initially declined at 10 weeks post tamoxifen but recovers by 20 weeks back to control levels, suggesting new OL genesis. To confirm this, we administered 5’-ethynyl-2’-deoxyuridine (EdU) in the drinking water to label dividing OPCs and their differentiated progeny (Supplementary Fig. 4a). We found few new (EdU+) OLs in Myrf^ΔiSox10^ mice, whereas there was elevated OL genesis in Myrf^ΔiPlp1^ mice that is sufficient to restore the total density of OLs to control levels by 20 weeks post tamoxifen (Fig. 2k, Supplementary Fig. 4c). The density of OLIG2+PDGFRα+ OPCs that incorporated EdU, as well as the total density of OLIG2+PDGFRα+ cells, increased in both Myrf^ΔiPlp1^ and Myrf^ΔiSox10^ relative to Myrf^fl/fl^ controls (Fig. 2h,j, Supplementary Fig. 4b), but did not differ between Myrf^ΔiPlp1^ and Myrf^ΔiSox10^ mice, nor did expression of transcripts indicative of proliferation (Supplementary Fig. 4e). These data indicate OPC proliferation is not altered in the absence of *Myrf*. Collectively, immunohistochemical and sequencing data demonstrate *Myrf* knockout from OL lineage cells in Myrf^ΔiSox10^ mice results in loss of OLs, with OPCs unable to differentiate beyond the COP level and generate new OLs.

### Remyelination failure is correlated with expansion of a microglia/macrophage population characterized by increased expression of lipid binding and metabolism transcripts

We next determined the influence of remyelination failure on the neuroinflammatory response. Demyelination in both Myrf^ΔiPlp1^ and Myrf^ΔiSox10^ mice is associated with an increase in microglia/macrophage density (Fig. 3a,b), astrogliosis (Supplementary Fig. 5a-e) and sparse T cell infiltration (Supplementary Fig. 6a). However, the densities of IBA1+ microglia/macrophages, and CD68 expression did not differ between Myrf^ΔiPlp1^ and Myrf^ΔiSox10^ mice (Fig. 3a-c). Densities of CD3+, CD3+CD4+, and CD3+CD8+ T cells also do not differ between Myrf^ΔiPlp1^ and Myrf^ΔiSox10^ mice (Supplementary Fig. 6b-e). Astrocytes in Myrf^ΔiSox10^ mice did show a marked upregulation of *Lcn2, Slc39a14,* and *Cp* transcripts, encoding proteins critical for the uptake and detoxification of iron (Supplementary Fig. 5e). We examined oxidative damage using an antibody for oxidized phosphatidylcholines but did not see an increase in Myrf^ΔiPlp1^ and Myrf^ΔiSox10^ mice (Supplementary Fig. 5f,g). Upregulation of these transcripts in astrocytes may, therefore, be an adaptive change to reduce oxidative stress in response to the loss of OLs, the major iron-storing cells of the CNS^26^.

**Fig. 3.**
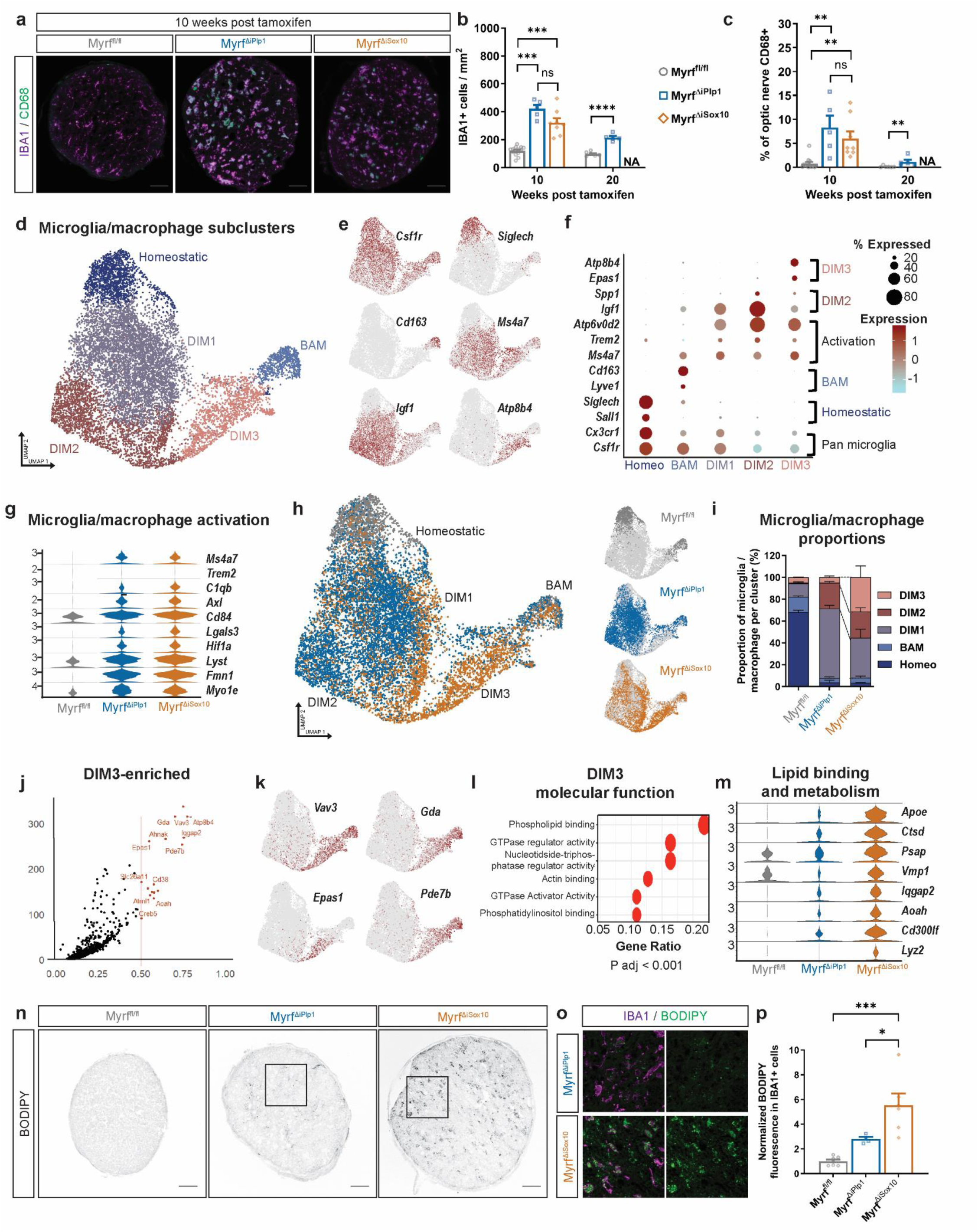
Remyelination failure in^ΔiSox10^ mice is associated with microglia/macrophages with elevated transcription of lipid metabolism genes and accumulated neutral lipids. **a** Optic nerve cross-sections stained with IBA1 for microglia/macrophages and the lysosomal marker CD68. **b** IBA1+ cells are increased in density in Myrf^ΔiPlp1^ (*P* = Q.QQQ5) and Myrf^ΔiSox10^ (*P* = 0.0010) relative to Myrf^fl/fl^ mice, but do not differ from each other (*P* = 0.1173) at 10 weeks post-tamoxifen in the optic nerve. IBA1 + microglia macrophages remain elevated in Myrf^ΔiPlp1^ over Myrf^fl/fl^ mice (*P* < 0.0001) at 20 weeks post tamoxifen, **c** The percent of the optic nerve occupied by CD63 staining is increased in in Myrf^ΔiPlp1^ (*P* = 0.0030) and Myrf^ΔiSox10^ (*P* = 0.0038) relative to Myrf^fl/fl^ mice, but do not differ from each other (*P* = 0.9999) at 10 weeks post-tamoxifen. At 20 weeks post-tamoxifen, the percent of the optic nerve that is CD68+ remains elevated in Myrf^ΔiPlp1^ relative to Myrf^fl/fl^ mice (*P* = 0.0087). **d** UMAP plot of reclustered microglia/macrophage nuclei identifying five annotated subclusters (homeostatic microglia, barrier associated macrophages (BAM), demyelination induced microglia/macrophages (DIM 1-3). *n* = 11,402 nuclei, **e** UMAP of key subcluster transcripts enriched within microglial/macrophage population; pan microglial/macrophage marker *(Csflr),* homeostatic microglia *(Siglech),* BAMs *(Cd163),* DIMs *(Ms4a7),* DIM2 *(Igf1)* and DIM3 *(Atp8b4).* **f** Dot plot showing expression of sub-cluster specific markers, **g** Violin plots for selected transcripts associated with activation in microglia/macrophage nuclei, **h** UMAP of microglia/macrophage lineage cell nuclei broken down by genotype, **i** Stacked bar graph of microglia/macrophage subcluster composition by genotype, **j** Volcano plot of key enriched transcripts in DIM3. Log-(fold change) > 0.5 adjusted p<0.05. Wilcoxon rank-sum test, **k** UMAPs of genes enriched in DIM3 cluster. **I** DIM3 gene ontology top six terms for molecular function, **m** Violin plots showing expression of select lipid binding and metabolism genes in the microglia/macrophage lineage by genotype, **n** BODIPY fluorescence in the optic nerve of Myrf^fl/fl^, Myrf^ΔiPlp1^’ and Myrf ^ΔiSox10^ mice at 10 weeks post tamoxifen, o Magnified region of the optic nerve from Myrf^ΔiPlp1^ and Myrf^ΔiSox10^ demonstrating the majority of BODIPY signal colabels with IBA1. **p** BODIPY fluorescence is increased in the optic nerve of Myrf^ΔiSox10^ mice relative to both Myrf^ΔiPlp1^ (*P* = 0.0258) and Myrf^fl/fl^ (*P* = 0.001). One-way Welch’s ANOVA with Dunnett’s T3 *post hoc* for week 10 in **b**, Kruskal Wallis with Dunn’s Test for 10 week comparison in **c** and one-way ANOVA with Tukey’s *post hoc* in **p**. For week 20 comparisons Student’s t-test was used in **b** and Mann Whitney U **c.** Error bars are SEM. Scale bars 50 pm in **a** and **n**. NA = Not applicable.

To more closely assess the microglial/macrophage response following remyelination failure in Myrf^ΔiSox10^ mice, we subclustered microglia from our snRNAseq dataset (Fig. 3d). We annotated five microglia clusters with three of these clusters (demyelination-induced microglia/macrophage 1-3; DIM1-3) enriched in both Myrf^ΔiPlp1^ and Myrf^ΔiSox10^ mice following demyelination. DIM1-3 are distinguished from homeostatic or barrier-associated macrophages (BAMs) by the presence of elevated activation markers, including *Ms4a7*, *Axl* and *Atp6v0d2* (Fig. 3e,f). The expression of these transcripts does not differ between Myrf^ΔiPlp1^ and Myrf^ΔiSox10^ mice, suggesting broadly similar activation (Fig. 3g). DIM2 microglia/macrophages are characterized by elevated levels of *Igf1* and *Spp1*, which have previously been found in a subpopulation of microglia along axon tracts^27^ (Fig. 3f). DIM3 is the only population that is increased in Myrf^ΔiSox10^ mice relative to the Myrf^ΔiPlp1^ line (Fig. 3h,i). This population is characterized by transcripts including *Vav3, Gda, Epas1 and Pde7b* (Fig. 3j,k). Gene set enrichment analysis on DIM3 enriched transcripts revealed the top term to be ‘phospholipid binding’ (Fig. 3l); accordingly, there is an upregulation of lipid binding and metabolism genes in Myrf^ΔiSox10^ microglia nuclei (Fig. 3m). When myelin is phagocytosed by microglia/macrophages, it undergoes lysosomal processing to produce cholesterol and fatty acids, which can be stored in lipid droplets and act as a reservoir for efflux^28^. BODIPY, a marker of neutral lipids present in lipid droplets^29^, accumulates in IBA1+ cells in Myrf^ΔiSox10^ relative to Myrf^ΔiPlp1^ (Fig. 3n-p). In summary, while both Myrf^ΔiPlp1^ and Myrf^ΔiSox10^ mice show similar microglia/macrophage numbers, a subset of microglia/macrophages in Myrf^ΔiSox10^ mice are characterized by upregulated lipid metabolism and binding transcripts and have increased lipid storage in the absence of remyelinating OLs.

### Remyelination is associated with protection from axonal degeneration and neuronal apoptosis

To determine how neuronal integrity is impacted during remyelination failure in Myrf^ΔiSox10^ mice, we assessed the health and integrity retinal ganglion cells (RGCs) and their axons in the optic nerve. There is an increase in the percentage of axons with accumulated organelles following demyelination in both Myrf^ΔiPlp1^ and Myrf^ΔiSox10^ mice at 10 weeks post tamoxifen, indicative of axonal transport deficits and injury^30, 31^ (Fig. 4a,b). By 20 weeks post tamoxifen in Myrf^ΔiPlp1^ mice, when remyelination is more complete, organelle accumulations are decreased relative to 10 weeks (Fig. 4b). Serum neurofilament light chain (NfL), a biomarker of axonal injury in a variety of diseases including MS^32^, is increased at 10 weeks post tamoxifen in both Myrf^ΔiPlp1^ and Myrf^ΔiSox10^ mice (Fig. 4c). Consistent with axonal accumulations, NfL levels fall in Myrf^ΔiPlp1^ mice by 20 weeks post tamoxifen and are indistinguishable from controls (Fig. 4c). Despite the presence of accumulations in both lines, quantification of total axonal numbers in the optic nerves reveals that detectable loss of axons relative to myelinated controls is only present in the Myrf^ΔiSox10^ line (Fig. 4d) and Myrf^ΔiPlp1^ mice do not feature axonal loss relative to controls at 10 or 20 weeks post tamoxifen.

**Fig. 4.**
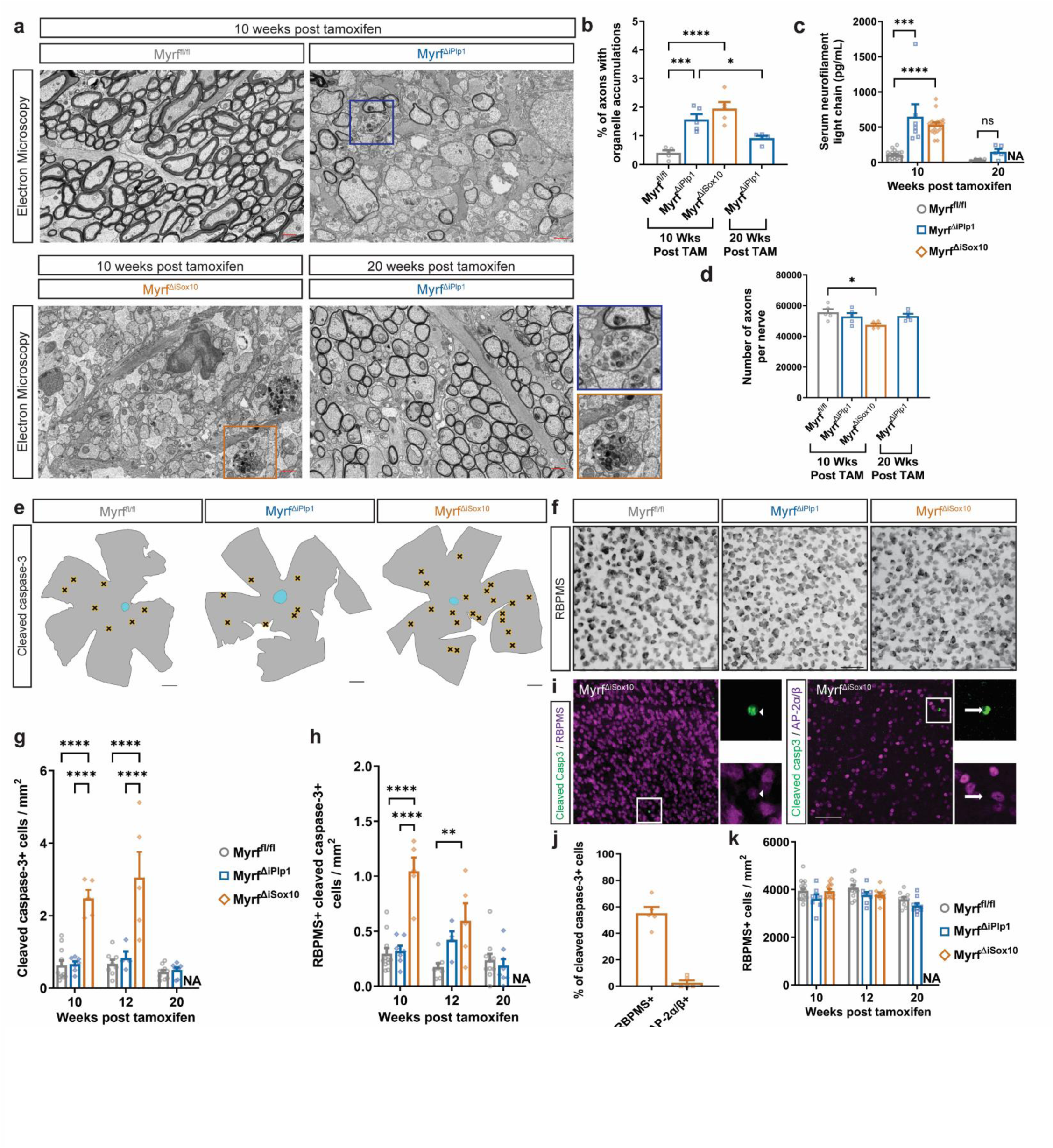
Myrf^ΔiSox10^ mice incur axon loss and have increased neuronal apoptosis relative to remyelinating Myrf^ΔiPlp1^ mice. **a** Electron micrographs of the optic nerve demonstrating axons with organelle accumulations in Myrf^ΔiPlp1^ and Myrf^ΔiSox10^ mice. Boxed areas show individual axons with organelle accumulation enlarged to the right, **b** The number of axons with organelle accumulations increase in Myrf^ΔiPlp1^*(P* = 0.0004) and Myrf^ΔiSox10^ mice *(P* < 0.0001) at 10 weeks post tamoxifen and falls in Myrf^ΔiPlp1^mice between 10 and 20 weeks post tamoxifen *(P* = 0.0437). **c** Serum neurofilament levels are increased at 10 weeks post tamoxifen in both Myrf^ΔiPlp1^ *(P* = 0.0004) and Myrf^ΔiSox10^ mice *(P* < 0.0001) but does not differ between Myrf^ΔiPlp1^ and Myrf^fl/fl^ mice at 20 weeks post tamoxifen (*P* = 0.1111). **d** The total number of axons in the optic nerve declines in Myrf^ΔiSox10^ mice relative to Myrf^fl/fl^ mice at 10 weeks post tamoxifen *(P* = 0.0202) but does not decline at 10 weeks post tamoxifen (*P* = 0.7086) or 20 post tamoxifen (*P* = 0.7765) in Myrf^ΔiPlp1^ relative to Myrf^fl/fl^ mice, **e** Overview of retina with locations of cleaved caspase-3+ cells in the GCL indicated by black X. **f** RBPMS in the retina across genotypes, **g** Increased density of cleaved caspase-3+ cells in Myrf^ΔiSox10^ relative to Myrf^fl/fl^ and Myrf^ΔiPlp1^ mice at 10 and 12 weeks post-tamoxifen (*P* < 0.0001 for referenced comparisons), **h** Density of cleaved caspase-3 and RBPMS double-labeled cells are increased in Myrf^ΔiSox10^ relative to Myrf^fl/fl^ and Myri^ΔiPlp1^at 10 weeks post tamoxifen (*P* < 0.0001) and increased relative to Myrf^fl/fl^ at 12 weeks post tamoxifen (*P* = 0.0018). **i** Co-labeling between cleaved caspase-3 and RBPMS in the retina. Arrowhead indicates cleaved-caspase-3+ cells colabeling with RBPMS. Arrow indicates cleaved caspase-3+ cell lacking AP-2a/p. **j** The majority of cleaved-caspase-3+ cells are RBPMS+ and few express AP-2a/p. **k** There is no loss of RBPMS RGCs in Myrf^ΔiPlp1^ or Myrf^ΔiSox10^ mice relative to Myrf^fl/fl^ controls (*P* >0.05 at all time points assessed). One-way ANOVA with Tukey’s *post hoc* for pairwise comparisons in **b**, and **d**. Kruskal Wallis with Dunn’s Test for comparisons at week 10 for **c**. For week 20, Student t-test used in **g** and **h** and Mann Whitney U for **c**. Two-ANOVA with Tukey’s *post hoc* for comparisons at week 10 and 12 in g and **h.** Error bars are SEM. Scale bars are 1 pm in **a**, 50 pm **f** and **i** and 500 pm **e**. NA = Not applicable.

We next asked whether the cell bodies of RGCs, which project their axons through the optic nerve, undergo apoptosis and loss in Myrf^ΔiSox10^ mice. Retinal flatmounts were stained with cleaved-caspase-3 to detect apoptotic cells (Fig. 4e) and with RBPMS (Fig. 4f), a pan RGC marker^33^. Myrf^ΔiSox10^ mice have increased apoptosis in the ganglion cell layer (GCL; Fig. 4g) and increased numbers of cleaved caspase-3+ RBPMS+ RGCs (Fig. 4h) relative to both the remyelinating Myrf^ΔiPlp1^ mice and non-demyelinated controls. We found that a portion of cleaved caspase-3 cells do not express RBPMS^34^, and we examined whether there is apoptosis in other cell types. Apoptotic cells are largely confined to the GCL, which consists of displaced amacrine cells and RGCs. Labelling amacrine cells with AP-2α and AP-2β, we found that only 2.7% ± 1.7% of the cleaved caspase-3+ cells are AP-2α+ or AP-2β+ in Myrf^ΔiSox10^ mice, whereas 55.2 ± 4.8% are RBPMS+ (Fig. 4j). RBPMS is downregulated during injury^34^, so it seems plausible that the remaining cleaved caspase-3+ cells represent late stage apoptotic RGCs in which the RBPMS protein has been lost (Fig. 4i). The apoptosis of RGCs cannot be attributed to *Myrf* knockout from neurons, as Sox10-CreERT does not recombine in RGCs and *Myrf* knockout within RGCs does not directly drive their apoptosis (Supplementary Fig. 7a-g). Neither Myrf^ΔiPlp1^ nor Myrf^ΔiSox10^ mice have a significant reduction of RGC density, likely owing to the rate of apoptosis in Myrf^ΔiSox10^ mice being insufficient to accrue a detectable loss over the time frame of these experiments (Fig. 4k). Taken together, both Myrf^ΔiSox10^ and Myrf^ΔiPlp1^ mice feature axonal swellings consistent with damage following demyelination. However, only Myrf^ΔiSox10^ mice, characterized by remyelination failure, culminate in axon loss relative to Myrf^fl/fl^ controls. Remyelination is associated with protection of RGCs from apoptotic cell death in Myrf^ΔiPlp1^.

### Increased activation of the DLK-mediated MAPKs and c-Jun in RGCs in Myrf^ΔiSox10^ mice with failed remyelination

To understand how chronically demyelinated neurons are damaged in Myrf^ΔiSox10^ mice, we performed bulk RNA-seq on the GCL (Fig. 5a). Laser dissecting the GCL greatly enriched for RGC-specific transcripts relative to the whole retina (Fig. 5b). The GCL of Myrf^ΔiSox10^ mice showed an upregulation of *Ecel1*, *Hrk,* and *Atf3* relative to Myrf^fl/fl^ (Fig. 5c). *In situ* hybridization of *Ecel1* and *Hrk* confirmed these transcripts are expressed in *Rbpms*+ RGCs in Myrf^ΔiSox10^ mice (Fig. 5d,e). Notably, these transcripts are known to be upregulated following DLK activation^35, 36^. DLK can mediate a retrograde signal that couples axonal injury to transcriptional changes in the nucleus, and often triggers subsequent apoptosis in the central nervous system^35, 36, 37^. DLK and the related leucine zipper-bearing kinase (LZK)^38, 39^ mediate this retrograde signal by activating downstream mitogen-activated protein kinases (MAPKs) including MAPK kinase 4/7 (MKK4/7) and c-Jun N-Terminal Kinase 2/3 (JNK2/3), resulting in the phosphorylation and activation of the transcription factor c-Jun^40, 41, 42, 43^ (Fig. 5l). Activation of this pathway has not previously been identified after demyelination.

**Fig. 5.**
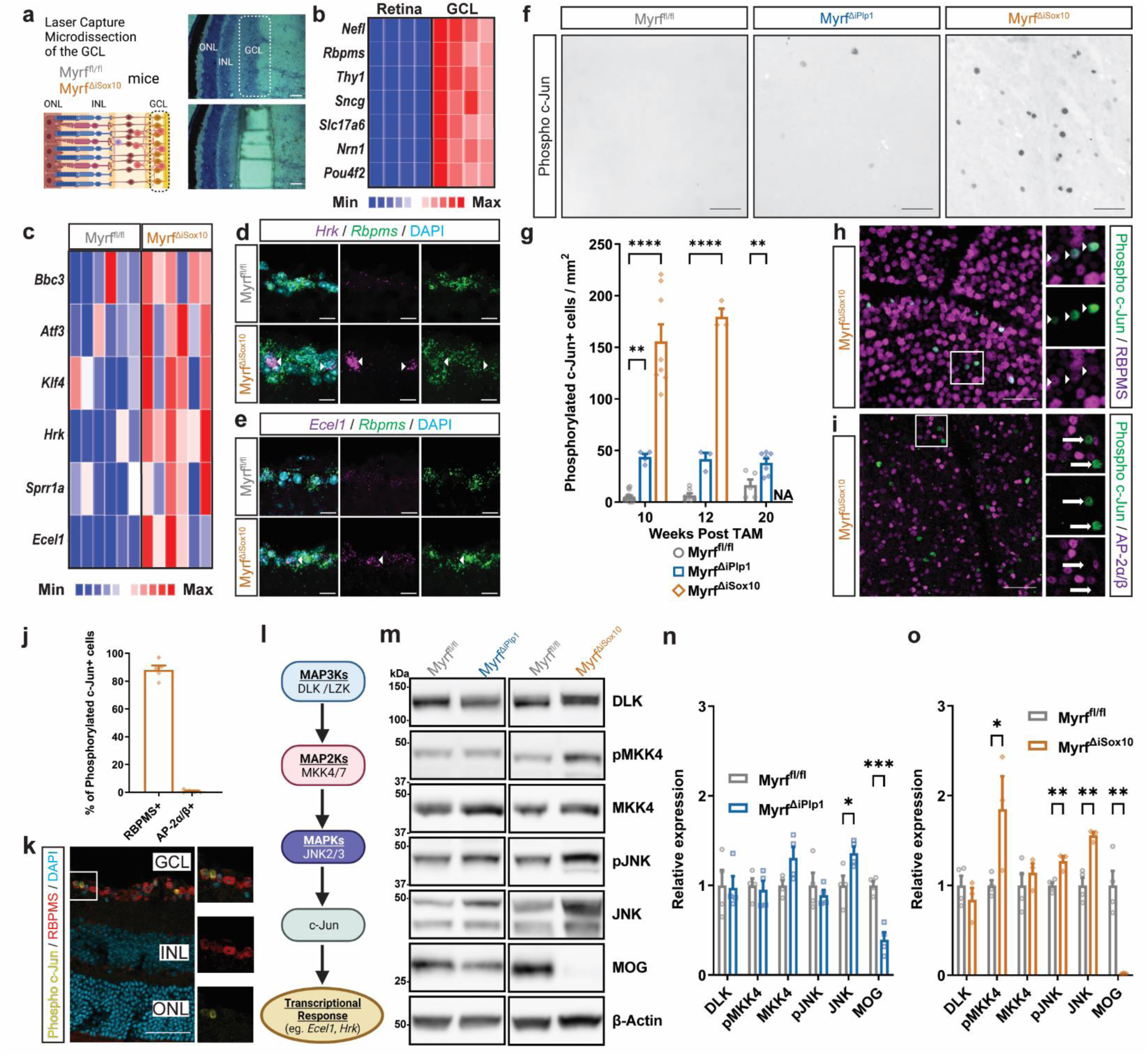
Activation of the DLK/JNK/c-Jun pathway in chronically demyelinated Myrf^ΔiSox10^ mice. **a** Experimental schematic and images of laser microdissection of the GCL of the retina, **b** Heatmap of expression of RGC-specific transcripts in the micro-dissected GCL samples relative to whole retina, **c** Heatmap of select transcripts from the GCL known to be activated by c-Jun/DLK signaling compared between genotypes, **d** *In situ* hybridization in the retina at 10 weeks post tamoxifen with probes against *Rbpms* and *Hrk.* Arrowheads indicate *Rbpms/Hrk* double-positive cells, e *In situ* hybridization in retina at 10 weeks post tamoxifen with probes against *Rbpms* and *Ecell.* Arrowheads indicate *Rbpms/Ecell* double-positive cells, **f** Retina at 10 weeks post tamoxifen stained with phosphorylated (Ser63) c-Jun. **g** Phosphorylated c-Jun is increased in Myrf^ΔiSox10^ mice at 10 and 12 weeks post tamoxifen relative to Myrf^fl/fl^ and Myf^ΔiPlp1^ *(P* < 0.0001 for all comparisons). The density of phosphorylated c-Jun+ cells is increased in Myrf^ΔiPlp1^ relative to Myrf^fl/fl^ at 10 and 20 weeks post tamoxifen (Wk10, *P* = 0.0088 and Wk20, *P* = 0.0092). **h** Example images of phosphorylated c-Jun within RBPMS+ cells in Myrf^ΔiSox10^ mice. Inlays are of boxed area. Arrowheads indicate colabeled cells. **I** Examples of phosphorylated c-Jun stained with AP-2a/p in Myrf^ΔiSox10^ retinae. Inlays are of boxed area. Arrows indicate phosphorylated c-Jun positive cells negative for AP-2α/β. **j** The vast majority of phosphorylated c-Jun+ cells are RBPMS+ and AP-2a/p-. **k** Image of retinal layers stained with phosphorylated c-Jun, RBPMS and DAPI. Inlays are of boxed area. **I** Schematic of the DLK-mediated MAPK cascade and c-Jun**. m** Western blot of optic nerves for DLK, pMKK4, MKK4, pJNK, JNK, MOG and P-actin loading control from optic nerves of Myrf^fl/fl^, Myrf^ΔiPlp1^ and Myrf^ΔiSox10^ mice, n Quantification of western blots. There is increased JNK levels (*P* = 0.0309) and decreased MOG in Myrf^ΔiPlp1^ (*P* = 0.0007) mice relative to Myrf^ΔiPlp1^ controls at 10 weeks post tamoxifen, o Quantification of western blots in Myrf^ΔiSox10^ mice relative to Myrf^fl/fl^. There is increased levels pMKK4 (*P* = 0.0434), pJNK (*P* = 0.0072), and total JNK (*P* = 0.0034) in Myrf^ΔiSox10^ mice at 10 weeks post tamoxifen. MOG levels are reduced in Myrf^ΔiSox10^ mice *(P* = 0.0038) relative to Myrf^fl/fl^ controls. Scale bars are 50 pm in **a f, h, i** and **k**, and 5pm in **d**, and **e**. Two-way ANOVA with Tukey’s *post hoc* at 10 and 12 weeks post tamoxifen, and Student’s t-test to compare groups at 20 weeks post tamoxifen in **g**. Student’s t-test in **n, o**. Error bars are SEM. NA = Not applicable.

To determine whether this pathway has elevated activation in the chronically demyelinated RGCs, we examined the phosphorylation of c-Jun in retinal flatmounts (Fig. 5f). The retinae of Myrf^ΔiSox10^ mice showed a large increase in the density of phosphorylated c-Jun+ cells relative to both remyelinating Myrf^ΔiPlp1^ and Myrf^fl/fl^ control mice (Fig. 5g). Nearly all these phosphorylated c-Jun+ cells co-labeled with RBPMS (Fig. 5h,j), with virtually no colocalization in AP-2α/β+ amacrine cells (Fig. 5i,j) and phosphorylation of c-Jun was largely confined to the GCL (Fig. 5k). In Myrf^ΔiSox10^ mice, the onset of c-Jun phosphorylation is coincident with decreased myelination and OL density at 8 weeks post tamoxifen, indicating a tight temporal relationship between demyelination and c-Jun phosphorylation in RGCs (Supplementary Fig. 8a-g). Inflammatory demyelination in experimental autoimmune encephalomyelitis (EAE) also results in the phosphorylation of c-Jun in RGCs (Supplementary Fig. 9a,c,d). Phosphorylation of c-Jun is not confined to RGCs, as spinal neurons also had elevated expression of phosphorylated c-Jun in Myrf^ΔiSox10^ mice (Supplementary Fig. 10b,d).

We next examined if MAPKs downstream of DLK are induced following remyelination failure in Myrf^ΔiSox10^ mice. Both the phosphorylation of JNK and MKK4 are increased in the optic nerves of Myrf^ΔiSox10^ mice at 10 weeks post tamoxifen (Fig. 5m,o), whereas neither kinase has detectable increases in phosphorylation in Myrf^ΔiPlp1^ mice (Fig. 5n). Although total DLK levels are not altered in Myrf^ΔiSox10^ mice relative to controls, the apparent molecular weight of the protein increases, consistent with post-translational modifications such as palmitoylation previously demonstrated after axonal injury^44, 45^. In summary, Myrf^ΔiSox10^ mice have increased phosphorylation of MAPKs, c-Jun and transcription of genes associated with axonal injury and stress.

### Pharmacological inhibition of DLK blocks apoptosis of demyelinated RGCs

To determine if DLK-mediated signaling is necessary to drive the activation of downstream c-Jun signaling and apoptosis in neurons following chronic demyelination, we treated demyelinated Myrf^ΔiSox10^ mice with the DLK inhibitor GNE-3511^46^ for three days via oral gavage (Fig. 6a). This regimen has been shown to be effective at blocking c-Jun phosphorylation after optic nerve crush^36^ and led to average serum levels over 4µM (Fig. 6b). At ten weeks post tamoxifen, retinal flatmounts were stained with phosphorylated c-Jun, or cleaved caspase-3 (Fig. 6c,d). Treatment of Myrf^ΔiSox10^ mice with GNE-3511 returned the phosphorylation of c-Jun to baseline levels observed in myelinated Myrf^fl/fl^ littermates (Fig. 6e). Likewise, cleaved-caspase-3+ cells and cleaved-caspase-3+RBPMS+ levels return to baseline, indicating near-complete protection from apoptosis of RGCs following acute GNE-3511 treatment (Fig. 6f,g). Thus, DLK inhibition completely blocks c-Jun phosphorylation and apoptosis of chronically demyelinated RGCs.

**Fig. 6.**
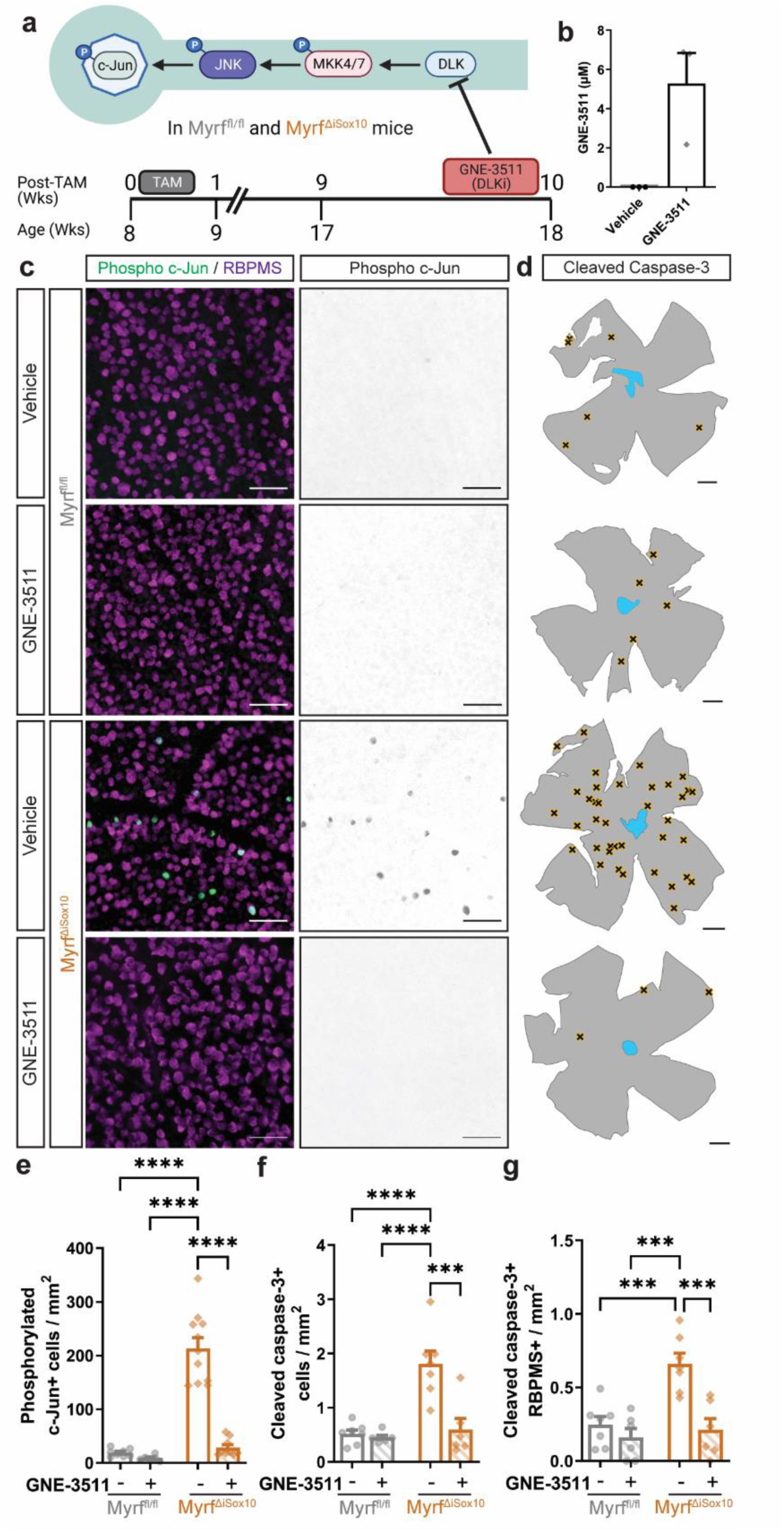
Pharmacological inhibition of DLK reduces c-Jun phosphorylation and blocks neuronal apoptosis in demyelinated Myrf^ΔiSox10^ mice. **a** Schematic of the DLK MAPK cascade and timeline of GNE-3511 administration, **b** Serum levels of GNE-3511 following 3 days of oral gavage, **c** Example retinal flatmount images of vehicle and GNE-3511 treated mice stained with phosphorylated c-Jun and RBPMS at 10 weeks post­tamoxifen. **d** Overview of retinas with locations of cleaved caspase-3+ cells in the ganglion cell layer (GCL) indicated by black X. **e** GNE-3511-treated Myrf^ΔiSox10^ mice show a significant reduction in the density of phosphorylated c-Jun positive RGCs relative to vehicle-treated Myrf^ΔiSox10^ and both vehicle-treated and GNE-3511-treated Myrf^fl/fl^ mice (*P* < 0 0001 for all comparisons) **f** GNE-3511-treatment of Myrf^ΔiSox10)^ mice at 10 weeks post-post tamoxifen leads to a reduction in cleaved caspase-3+ cells in the GCL relative to vehicle-treated Myrf^ΔiSox10^ controls tamoxifen (*P* = 0.0002) and Myrf^fl/fl^ mice (*P* < 0.0001). g Cleaved caspase-3+ RBPMS+ cells are decreased in GNE-3511-treated Myrf^ΔiSox10^ relative to vehicle-treated mice at 10 weeks post-tarn oxifen (*P* = 0.0005) and relative to My controls (vehicle, *P*= 0.0001, GNE-3511, *P* = 0.0008). Two-way ANOVA with Tukey’s post hoc for individual pairwise comparisons in **f, g** and **h**. Error bars are SEM. Scale bars are 500 pm in **d** and 50 pm in **c**.

### Genetic disruption of DLK inhibits apoptosis of demyelinated RGCs

We next used a CRISPR/Cas9 genetic disruption approach to determine the contributions of DLK, and its related kinase LZK, to apoptosis of chronically demyelinated RGCs. DLK and LZK can have partially redundant roles^38^, so we developed sgRNAs to disrupt one or both genes within RGCs (Fig. 7a). We injected adeno-associated viruses (AAVs) containing sgRNAs targeting either DLK (*Map3k12*), LZK (*Map3k13*) or both DLK and LZK intravitreally at six weeks post tamoxifen in Myrf^ΔiSox10^ and Myrf^fl/fl^ mice crossed with mice constitutively-expressing Cas9 (Myrf^ΔiSox10^ Cas9 or Myrf^fl/fl^ Cas9). Control AAVs with sgRNAs targeting eGFP and LacZ were injected into the contralateral eye. All AAVs effectively labeled RGCs throughout the retina (Fig. 7b). To determine whether Cas9-mediated disruption of DLK and/or LZK prevented the phosphorylation of c-Jun and apoptosis of RGCs, we stained retinal flatmounts with phosphorylated c-Jun (Fig. 7c) and cleaved caspase-3 (Fig. 7d). Retinae infected with either sgDLK alone or sgDLK/sgLZK have near-complete suppression of both c-Jun phosphorylation and RGC apoptosis (Fig. 7e,g,h,j). In contrast, knockout of LZK alone had a no significant effect on either phosphorylated c-Jun or cleaved caspase-3 cell density (Fig 7f,i). Together, these data indicate DLK is the major MAP3K necessary for neuronal apoptosis following demyelination and subsequent remyelination failure.

**Fig. 7.**
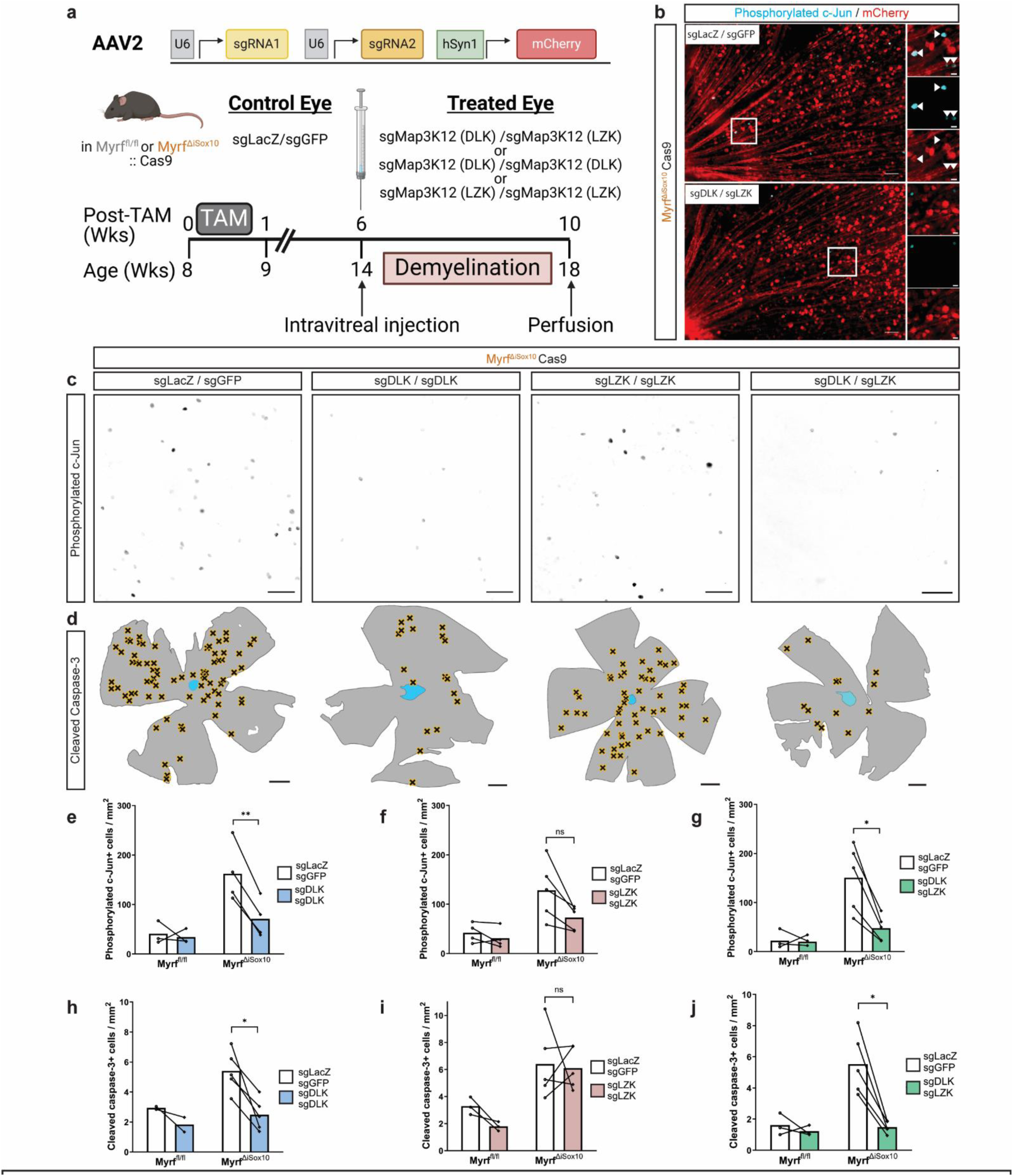
DLK is necessary for neuronal apoptosis following demyelination in Myrf^ΔiSox10^ mice. **a** Schematic of the viral CRISPR/Cas9 approach to disrupt DLK and/or LZK in retinal cells, **b** Retinae stained with phosphorylated c-Jun and mCherry at 10 weeks post tamoxifen following treatment with sgLacZ/sgGFP, or sgDLK/sgLZK on the opposite eye. Inlays are of boxed areas and scale bar is 5 pm in boxed areas, c Representative images of phosphorylated c-Jun immunostaining in retinal flatmounts following administration of sgLacZ/sgGFP. sgDLK/sgDLK, sgLZK/sgLZK, sgDLK/sgLZK. **d** Overview of retinas with locations of cleaved caspase-3+ cells in the GCL indicated by black Xs in viral-treated eyes, **e** Reduced density of phosphorylated c-Jun cells within the GCL following sgDLK/sgDLK administration in Myrf^ΔiSox10^ mice (*P* = 0.0080). **f** No change in the number of phosphorylated c-Jun positive cells in the GCL following sgLZK/sgLZK administration (*P* = 0.0766). **g** Reduced phosphorylated c-Jun cells within the GCL following sgDLK/sgLZK administration in Myrf^ΔiSox10^ mice (*P* = 0.0148). **h** Reduced cleaved-caspase-3+ cells within the GCL following sgDLK/sgDLK administration in Myrf^ΔiSox10^ mice (*P* = 0.0254). i No change in the number of cleaved-caspase-3+ cells within the GCL after sgLZK/sgLZK administration (*P* = 0.8530). **j** Reduced cleaved-caspase-3+ cells within the GCL following sgDLK/sgLZK administration in Myrf^ΔiSox10^ mice (*P* = 0.0125). Scale bars are 500 pm In **d** and 50 pm in band **c**. Connected lines indicate retinae from the same mouse in **e-j**. Paired student’s t-test with Holm-Sidak correction for multiple comparisons was used from **e-j**. ns = not statistically significant. Error bars are SEM.

## Discussion

By comparing our remyelination capable and deficient mouse lines, we find remyelination is associated with improved functional recovery and neuronal integrity. Mice with remyelination failure have elevated activation of MAPKs downstream of DLK and phosphorylation of the transcription factor c-Jun. We demonstrate that DLK is necessary for apoptosis of demyelinated RGCs using Cas9-mediated disruption of DLK from RGCs and pharmacological inhibition of DLK, both sufficient to suppress c-Jun phosphorylation and RGC apoptosis. We propose that neuroprotection in demyelinating disease can be achieved by targeting retrograde signaling mediated by DLK within demyelinated neurons or by promoting remyelination which is associated with suppression of the same DLK-mediated signaling cascade (Supplementary Fig. 11).

The two models of genetic demyelination induced by knockout of *Myrf* offer distinct advantages for subsequent studies. Myrf^ΔiPlp1^ mice, which knock *Myrf* out from OLs, demonstrate highly reproducible, CNS-wide demyelination with clear behavioral and physiological readouts. The remyelination process begins rapidly in Myrf^ΔiPlp1^ mice, even before all myelin destined to be lost has degenerated. We found the delay of conduction within the visual system correlates temporally with demyelination, whereas restoration of axonal conduction is associated with remyelination in Myrf^ΔiPlp1^ mice. Likewise, motor function declines significantly with the onset of demyelination and slowly recovers during remyelination, as seen in other models of extensive demyelination^47,48^. In contrast to Myrf^ΔiPlp1^ mice, Myrf^ΔiSox10^ mice, do not effectively remyelinate and fail to recover, with the differentiation of remyelinating cells blocked at the COP stage. Myrf^ΔiSox10^ mice are the first model that cell-selectively induces demyelination and impairs remyelination, allowing for investigation of how chronic demyelination impacts neurons and the efficacy of neuroprotective therapeutics.

We use these models to determine the degree of neurodegeneration in mice without effective remyelination relative to those mice with efficient remyelination. An association between remyelination and neuroprotection has been difficult to establish in rodents, in part due to their rapid remyelination. Targeting the OL lineage with cell-specificity, we find greater RGC apoptosis in mice Myrf^ΔiSox10^ mice which are unable to remyelinate. Apoptosis of RGCs is persistent at 10- and 12-weeks post tamoxifen in the Myrf^ΔiSox10^ mice, supporting a notion of continuous neurodegeneration in the context of prolonged demyelination. Although the rate of apoptosis in RGCs is relatively modest (∼0.06% of RGCs being apoptotic at any given time), the cumulative effect of such apoptosis on the population would be significant over time given the rapid rate of microglial clearance of apoptotic cells. Accelerating remyelination by deleting the M1 muscarinic receptor from OL lineage cells increases axon preservation and functional recovery in EAE^49^. Collectively, this lineage-specific gain of function experiment and our loss of function experiments provide compelling support that remyelination rate is a critical determinant of subsequent neurodegeneration. Additionally, both neuroimaging^50, 51, 52^ and histopathological^14, 53^ studies measuring remyelination in MS lesions support a neuroprotective role. One limitation is our approach is incapable of dissociating the relative neuroprotective contributions of new OLs verses the deposition of new myelin. Future studies targeting specific mechanisms by which OLs may support neuronal health during remyelination will be crucial to disentangle the role of the oligodendrogenesis versus myelinogenesis on alleviating neurodegeneration.

Our experiments provide an important contrast to recent findings that myelinated axons are more at risk for degeneration during autoimmune-mediated demyelination^30^ and in *Plp1* mutants^54, 55^. Axonal damage in EAE is more prevalent in myelinated axons relative to their demyelinated counterparts, and hypomyelinated *Mbp* mutant mice incur less axonal damage during EAE^30^. Plp1 over-expressing mice, mimicking *Plp1* duplication in humans, initiate progressive demyelination during early adulthood^55^. In these mice, axons are most vulnerable to damage during active demyelination and are relatively protected following the resolution of inflammation^55^. Likewise, *Plp1* mutants with abnormal myelin are relatively protected from CD8+ T cell-mediated neurodegeneration when demyelinated^54^. Both in MS tissue^14, 53^ and in experimental models^30, 55^, chronically demyelinated neurons seem less vulnerable to damage relative to those undergoing active demyelination. Thus, axons are highly vulnerable during active demyelination or when deposited with dysfunctional myelin. Nevertheless, in Myrf^ΔiSox10^ mice following chronic demyelination apoptotic RGCs are found at a more than four-fold increase relative to remyelinated and control mice, suggesting remyelination failure is associated with slow, but persistent neurodegeneration. Collectively, these animal studies indicate a substantial challenge for targeting myelin in MS; during the early stages of the disease damaged myelin is harmful to the axon and needs to be cleared and inflammation resolved, yet prolonged demyelination is also detrimental to both the function and viability of the neuron.

An important limitation of our studies is it remains unclear the contribution of inflammation to neurodegeneration following remyelination failure in Myrf^ΔiSox10^ mice. Microglia have a crucial role in regulating the efficacy of remyelination^56, 57, 58^. Interestingly, we found the reciprocal is also true: preventing oligodendrogenesis and remyelination in Myrf^ΔiSox10^ mice led to expansion of a microglial population, which could conceivably damage neurons. This microglial population is characterized by the elevated expression of lipid-binding and metabolism transcripts. Cholesterol is the major lipid constituent of myelin and both sterol synthesis and efflux in microglia are necessary for effective remyelination^59, 60^, presumably by acting as a source of sterols for newly generated OLs. In Myrf^ΔiSox10^ mice, it is intriguing to consider that in the absence of new remyelinating OLs, microglia lack their normal cellular sink for recycled cholesterol from damaged myelin. In support of impaired cholesterol export from microglia, we found an accumulation of neutral lipid droplets, which act to store cholesterol esters in microglia from Myrf^ΔiSox10^ mice. Increased accumulation of cholesterol within microglia and poor efflux is associated with persistent inflammation^60, 61^. With disease chronicity in MS, microglia become more diffusely activated^62^; it is conceivable that poor remyelination may contribute to persistent microglial activation and potentially neurodegeneration. Unfortunately, given the premature death of Myrf^ΔiSox10^ mice, we cannot determine whether remyelination failure would continue to result in prolonged microglial inflammation and if that would subsequently damage neurons. Future work should determine if this this shift in microglia phenotype following remyelination failure directly contributes to neurodegeneration.

While remyelination failure has been associated with axonal damage in MS^14, 53^, it has remained unclear how demyelination-induced damage to the axon may be discerned in the nucleus or if it drives loss of the soma. Periventricular lesions that fail to remyelinate in MS are associated with greater atrophy of brain regions they project from^51^, suggesting that demyelination of the axon may be perceived by the soma and culminate in neurodegeneration. Here, we find chronic demyelination of the axon is associated with elevated DLK-mediated phosphorylation of the transcription factor c-Jun in the nucleus, transcriptional changes and RGC apoptosis. Potentially, DLK could be activated in Myrf^ΔiSox10^ mice either as a direct loss of OL support or due to other factors such as altered inflammation. However, DLK is localized within the axon and several stimuli known to drive DLK-mediated signaling including axonal cytoskeletal disruption^63^ as well as transport deficits^64^, were observed following demyelination. Mechanistically, injury to the axon results in palmitoylation of DLK and its localization to retrogradely trafficked vesicles; a process required for phosphorylation of downstream kinases including JNK^45^. JNK signaling is a well-known activator of c-Jun mediated transcription in response to axonal injury^35, 37^, generally triggering apoptosis of CNS neurons^65^. c-Jun likely regulates apoptosis at least in part through modulating the expression of *Bcl2* family members like *Bcl2l2* and *Bbc3*^65, 66^. Taken together, we suggest DLK mediates a retrograde signal from the demyelinated axon that culminates in phosphorylation of c-Jun and subsequent apoptosis.

We found that DLK inhibitors protect neurons from apoptosis following chronic demyelination. Notably, we also see c-Jun phosphorylation and RGC apoptosis following inflammatory demyelination in EAE. This suggests the pathway may also be induced during acute inflammatory demyelination and that DLK inhibitors may also be protective in this context. Blocking DLK activity is therefore a strong candidate for neuroprotection in demyelinating disease. While a number of blood-brain barrier permeable DLK inhibitors have already been developed^46, 67^, a recent phase I trial with a DLK inhibitor in ALS warrants caution as prolonged DLK inhibition was associated with considerable safety concerns^68^. It may be necessary to dissect and target the downstream responses induced by DLK that drive neurodegeneration in the context of remyelination failure to produce druggable therapeutic targets. However, our data strongly supports that remyelination is also capable of suppressing DLK-mediated signaling and neuronal apoptosis. Even the incomplete remyelination present within the optic nerve of Myrf^ΔiPlp1^ mice at 10 weeks post tamoxifen is associated with suppressed MAPK signaling and apoptosis relative to the Myrf^ΔiSox10^ line. Therefore, it is plausible that even promoting a relatively small degree of remyelination will reduce neuronal attrition and will be of value in treating demyelinating disease.

## Acknowledgments

We would like to thank Brett Hilton for critically reading and providing feedback on this manuscript. We thank Pamela Canaday and Christina Metea for their assistance and guidance with flow cytometry. Alex Klug and Amy Carlos provided 10x microfluidics and library production support. Brian Jenkins, Hannah Bronstein and Stefanie Kaech Petrie provided guidance for microscopy image analysis. Paul Meraner kindly provided Cas9-expressing Neuro2a cells for sgRNA validation. This work received support from the Flow Cytometry, Gene Profiling/RNA and DNA Services Shared Resource, OHSU Molecular Virology Core, Massively Parallel Sequencing Shared Resource, Multiscale Microscopy Core (MMC) and Advanced Light Microscopy Cores at OHSU. This work was supported by grants from Race to Erase MS, the NINDS (R01NS120981), NINDS P30 (NS061800), Collins Medical Trust (1016373), the National Multiple Sclerosis Society (RG-2001-35775), American Heart Association (20CDA35320169) and National Institute of Diabetes and Digestive and Kidney Diseases (K01DK121737). G.J.D. was supported by a postdoctoral fellowship (FG-1808-32238) and a career transition award (TA-2105-37636) from the National Multiple Sclerosis Society. C.C. was supported by FISM - Fondazione Italiana Sclerosi Multipla – cod. 2019/BC/002 and financed or co-financed with the ‘5 per mille’ public funding. B.E. was supported by an endowment from the Warren family. G.J.D. and B.E. thank Caron and Larry Ogg for their generous financial support to support G.J.D and helping to make this work possible. All schematics created with BioRender.com were exported on a paid subscription with publication licenses.

## Contributions

G.J.D. and B.E. designed and conceived the experiments. G.J.D., S.D.I., J.H., M.M., J.E.J., C.C., A.A, N.J., K.A., S.J.F., B. Stedelin, B. Sivyer, and B.E. performed experiments and acquired data. G.J.D., K.E. and J.N. curated data and provided code for analysis of the snRNAseq data. G.J.D, K.E., C.C., A.A., J.N. and B.E. analyzed data. A.M. trained and supervised G.J.D. in acquisition of CAPs from the optic nerve. J.N. trained and supervised G.J.D. and K.E. in snRNAseq analysis. T.A.W. provided expertise with MAPK reagents and biology. S.A.A., A.M., T.S., A.J.G., and B.E. provided resources or funding for the experiments. G.J.D. and B.E. wrote the manuscript with inputs from all authors. B.E. supervised the project.

## Methods

### Mouse lines and husbandry

All mice were housed and maintained in the Oregon Health & Science University animal facility in a pathogen-free temperature and humidity-controlled environment on a 12-hour light/dark cycle. All animal procedures were performed in accordance with, and approved by, the Institutional Animal Care and Use Committee of OHSU. Myrf^fl/fl^ mice were generated in the Barres laboratory^15^ (B6;129-*Myrf^tm1Barr^*/J, JAX: 010607). After being crossed to C57BL/6 for over ten generations they were crossed to either Plp1-CreERT mice^69^ (B6.Cg-Tg[Plp1-cre/ERT]3Pop/J, JAX:005975) or Sox10-CreERT mice^19^ (CBA;B6-Tg[Sox10-icre/ERT2]388Wdr/J, JAX:026651). To allow for CRISPR/Cas9 mediated disruption of genes, Myrf^ΔiSox10^ mice were crossed to constitutively-expressing Cas9 mice^70^ (Gt[ROSA]26Sortm1.1[CAG-cas9*,-EGFP]Fezh/J, JAX:024858) to produce Myrf^ΔiSox10^ Cas9 and Myrf^fl/fl^ Cas9 mice. In all cases CreERT negative littermates served as non-demyelinated controls, and were matched by sex to knockout mice when possible. Plp1-CreERT mice and Sox10-CreERT mice were crossed with inducible Sun1-GFP mice^71^ (B6.129-Gt(ROSA)26Sortm5.1(CAG-Sun1/sfGFP)Nat/MmbeJ, JAX 030952) to assess which cells are recombined within the retina. Genotypes were determined by PCR analysis of ear clips, using established primers for each line, and revalidated at experimental endpoint. All experiments were conducted in both sexes of eight week-old mice. Every week following tamoxifen administration mice were weighed and health assessments performed. When mice reached a motor score of three, indicated by ataxia and significant hindlimb weakness, they received wet food on the bottom of their cage to maintain hydration. Initial cohorts indicated that Myrf^ΔiSox10^ mice developed seizures after 12 weeks post tamoxifen, so all analyses were limited to 12 weeks or earlier.

### Tamoxifen administration

Tamoxifen (T5648, Sigma) was dissolved in corn oil (C8267, Sigma) at 20 mg/mL using heat (37°C) and agitation. Mice received intraperitoneal injections at 100 mg/kg for five consecutive days at eight weeks of age. Tamoxifen was prepared fresh prior to administration for each cohort of mice.

### EdU administration

To examine proliferation of OPCs and differentiation of new OLs, we administered 5-ethynyl-2′-deoxyuridine (EdU) in the drinking water starting the week after tamoxifen injections until 10 weeks post-tamoxifen. EdU (NE08701, Carbosynth) was dissolved in water along with 0.2 mg/mL of dextrose (D16-500, Fisher) to encourage consumption. EdU water was changed every two to three days over the course of administration.

### GNE-3511 administration

GNE-3511 (5331680001, Millipore-Sigma) was emulsified into 0.5% methylcellulose (M7140, Sigma) with 0.2% Tween 80, and vortexed prior to oral gavage^36^. At 10 weeks post tamoxifen administration, Myrf^ΔiSox10^ mice and Myrf^fl/fl^ controls received two daily gavages with either GNE-3511 at 75 mg/kg or vehicle only. Gavages were at least eight hours apart for three consecutive days, for a total of six gavages.

### EAE induction

Experimental autoimmune encephalomyelitis (EAE) was induced in eight week-old female C57BL/6 mice via immunization with myelin oligodendrocyte glycoprotein (MOG)_35-55_ (PolyPeptide Laboratories). 200 µg MOG_35-55_ was dissolved in complete Freund’s adjuvant containing 400 µg of *Mycobacterium tuberculosis* (231141, Difco) and injected subcutaneously in 0.2 mL volume per mouse. Pertussis toxin (181, List Biological labs Inc) was administered via intraperitoneal injection after the MOG_35-55_ injection and two days later at 75 ng and 200 ng per mouse doses, respectively. Mice were perfused within three days of peak disease and all mice reached an EAE score of three. Following perfusion, retinae were prepared for flatmount staining and optic nerves frozen and subsequently cryosectioned.

### Intravitreal injections

Intravitreal injections of viruses containing tandem sgRNAs were used to induce Cas9-mediated disruption of *Map3k12* and *Map3k13.* Myrf^fl/fl^ Cas9 and Myrf^ΔiSox10^ Cas9 mice were anaesthetized with ketamine/xylazine (ketamine 100 mg/kg and xylazine 12 mg/kg). Proparacaine (NDC 13985-611-15, Akorn Inc) and tropicamide (NDC 17478-102-12, Akorn Inc) eye drops were applied to provide local analgesia and better visualization of the injection, respectively. One eye received AAV expressing small guide RNAs against LacZ and GFP under U6 promoters (AAV2-U6-sgLacZ-U6-sgGFP-hSyn1-mCherry) with the opposite eye receiving sgRNAs against *Map3K12* (DLK), *Map3K13* (LZK), or both genes. The concentration of viruses was adjusted to 2.8 x10^12^ genome copies per mL immediately prior to injection in sterile 1x phosphate buffered saline (PBS). Under stereo microscopic control, the ora serata was incised with a 30 gauge needle without touching the lens. AAV vectors (1 µL/injection) were delivered into the vitreous through the incision using a 5 µL Hamilton microinjection syringe with a blunt 20 deg 33 gauge needle. After injection, paralube ophthalmic ointment (NDC 17033-211-38, Dechra) was applied and mice were placed on a heating pad until they awoke.

To determine if *Myrf* knockout of from RGCs induces apoptosis, AAV2-hSyn1-Cre along with AAV2-hSyn1-mCherry (Addgene114472-AAV2) were mixed at 2.32 x 10^12^ genome copies per mL prior to injection and intravitreally injected into one eye of Myrf^fl/fl^ mice. The control eye received just AAV2-hSyn1-mCherry. These injections were conducted as above, and mice were perfused 10 weeks and retinae harvested for flatmount staining.

### Optic Nerve Crush

Mice were prepared for surgery with the same ketamine/xylazine protocol for anesthesia as above. The right eye of each animal received an incision of the superior conjunctiva exposing the optic nerve, with care to avoid lesioning the orbital sinus. The nerve was crushed approximately 0.5 mm from the optic disk for 10 seconds using fine forceps (Dumont, #5). Following surgery paralube ophthalmic ointment was applied and 3.25 mg/kg of extended-release buprenorphine (Ethiqa XR formulation) was administered subcutaneously. The mouse was placed on a heating pad until it was awake.

### Compound action potential recordings

Mice were deeply anaesthetized with ketamine and xylazine as above and an incision was made to expose the dorsal skull. The optic nerves were cut just behind the optic disk. The dorsal skull was then removed and olfactory bulbs were cut with fine surgical scissors, and the brain was gently lifted to expose the optic chiasm. The optic chiasm was then cut and optic nerves were carefully removed and placed in oxygenated artificial cerebrospinal fluid (aCSF) (124mM NaCl, 1mM NaH_2_PO_4_, 2.5 mM KCl, 26 mM NaHCO_3_, 10mM glucose, 1 mM Na Ascorbate, 1mM MgCl_2_, 2mM CaCl_2_). Glass suction electrodes were prepared by heating the ends of glass capillaries over a flame until the tip was slightly constricted to match the nerve diameter, a few millimeters away the electrodes were bent at approximately 30 degrees to facilitate nerve holding. A silver wire was inserted into both recording and stimulating electrodes, and a reference wire was coiled around the electrodes to form a connection with the bath. Both electrodes were filled with aCSF. The optic nerve was transferred to a recording chamber continuously perfused with aCSF and held in place with wettened filter paper. Using gentle suction, the chiasmal side of the nerve was drawn into the recording electrode and the retinal side into the stimulating electrode. The nerve was then allowed to acclimate for 30 minutes to allow the tissue to form a seal with the constricted tip of the glass electrodes before recording began. The stimulating electrode was connected to a constant current isolated stimulator unit (Digitimer DS3) and nerves were stimulated at increasing amplitudes from 0.2mA to 2mA at 0.2 Hz for 100 µs until supramaximal threshold was found. Nerves were stimulated at 125% supramaximal threshold for recordings used in quantifications. Signals from the recording electrode were digitized via a Digidata 1440B, amplified using an Axon Instruments Multiclamp 700B amplifier and recorded using Clampex 10.7 software (Molecular Devices). CAP curves were subtracted from recordings following administration of TTX (1µm, HB1035 HelloBio). Data was then analyzed using the Clampfit10 software and to find both the latency to the highest peak and CAP area. CAP area is proportional to the number of stimulated axons^72^, and was measured as the area of the positive voltage deflection following stimulus. Each mouse was considered a biological replicate, and one nerve was measured from each mouse.

### Visual evoked potential recordings

Low ambient light (13-18 Lux) was ensured in the room where VEPs were recorded. Mice were anesthetized via intraperitoneal injection using both xylazine (12.5 mg/kg) and ketamine (87.5 mg/kg) diluted in PBS. Pupils were dilated with Tropicamide eye drops administered to each eye precisely six minutes after ketamine/xylazine administration. Mice were then placed in a sealed cardboard box (12×12×16 cm) for five minutes for dark adaptation. For the flash VEP one centimeter steel needle electrodes (Natus Neurology) were placed medially under the skin between the two eyes along the sagittal suture, with the needle inserted eight mm to optimize proximity to the visual cortex. A needle electrode inserted subcutaneously just above the tip of the nose served as reference electrode and an additional electrode inserted into the tail served as the ground electrode. A dome was next lowered to reduce ambient light and VEP stimulation and recording began precisely 13 minutes after administration of anesthesia. Flash based binocular visual electrophysiology was measured using an Espion Diagnosys system (Diagnosys LLC). Examinations consisted of three runs, with the following characteristics: pulse intensity 3 cd.s/m2, frequency 1 Hz, on-time 4 ms, pulse color: white-6500K, and 100 sweeps per acquisition as per^22^. The standard VEP waveform with these parameters was characterized by a prominent negative deflection after approximately 70 ms which we identified as N1. N1 was defined as the first negative deflection after 50 ms. The two most representative/reproducible waves were used for analysis. The exam was performed by an operator blinded to mouse genotype.

### Tissue Processing

Mice were deeply anaesthetized with ketamine (400 mg/kg) and xylazine (60 mg/kg) then transcardially perfused with 10 mL of PBS and 40mL of freshly hydrolyzed 4% paraformaldehyde (19210, Electron Microscopy Sciences). Tissues were gently dissected and placed in 4% paraformaldehyde for post fixation. For optic nerves used for electron microscopy, nerves were post fixed in 2% paraformaldehyde (15710, Electron Microscopy Sciences) with 2% glutaraldehyde (16310, Electron Microscopy Sciences) instead of 4% paraformaldehyde. For immunohistochemistry, optic nerves were post fixed for two hours, brains overnight, retinae for one hour and spinal cords for four hours. Optic nerves, brains, and spinal cords were then cryoprotected in 30% sucrose for at least 48 hours. Tissues were embedded in OCT and frozen on dry ice and stored at -80°C until sectioning on a cryostat (CM3050-S, Leica). Brain sections were mounted at 10 µm thickness on Superfrost Plus slides (1255015, Fisher Scientific) between 1.4mm to -0.6mm relative to bregma. Optic nerve sections were made between 2-4mm retinal to the optic chiasm mounted at 10 µm thickness. All sections were stored at -80°C until immunohistochemistry was performed. Eyes were washed with 1x PBS following post-fixation and the retinae were dissected and post-fixed in 4% paraformaldehyde for 30 minutes and washed three times in 1x PBS. Retinae for *in situ* hybridization were flash frozen on dry ice following intracardiac perfusion with 1x PBS.

### Immunohistochemistry

Slides were thawed from the -80°C until dry and then rehydrated in 1x PBS. For MBP staining, tissue delipidization was performed by immersing slides in ascending and descending ethanol solutions before being washed 3x in 1x PBS. Slides were blocked for 30 minutes at room temperature with 10% fetal calf serum (SH30910.03, Cytiva) with 0.2% Triton-X100 (10789704001, Sigma). Primary antibodies were applied overnight in 1x PBS with 0.2% Triton-X100 in a sealed container at room temperature. Primary antibodies included mouse anti-BCAS1 (1:200; sc-136342, Santa Cruz), chicken anti-MBP (1:200; MBP, Aves), mouse anti-CC1 (1:500; OP80, Millipore), rabbit anti CD3 (1:500; NB600-1441SS, Novus), rat anti CD4 (1:200; MAB554, R and D Systems), rat anti CD8 (1:200, 550281, BD Biosciences), rabbit anti-OLIG2 (1:500; AB9610, Millipore), chicken anti OLIG2 (1:500, OLIG2-002, Aves), goat anti mouse-PDGFRα (1:200 AF1062, R&D Systems), rat anti-CD68 (1:500; MCA1957GA, Biorad), rabbit anti-Iba1 (1:1000; 019-19741, Wako), rabbit anti SYT4 (1:200; 105 143 Sysy Antibodies), mouse anti NKX2.2 (1:200, 74.5A5 DSHB), rabbit anti-GFAP (1:1000; Z0334, Dako), mouse anti NFL-DegenoTag (1:1000, MCA-6H63, Encor Biotechnology), mouse anti-E06 (Oxidized phospholipid, 1:500, 330001S, Avanti), chicken anti-mCherry (1:1000, mCherry-0100, Aves), chicken anti-GFP (1:1000, ab13970, Abcam), rabbit anti-cleaved caspase-3 (1:200; 559565, BD Pharminogen) and rabbit anti-cleaved caspase-3 (1:200; AF835, R and D Systems). Following incubation with primary antibodies, slides were washed three times in 1x PBS before appropriate Alexa Fluor 488, 555 or 647 secondary antibodies (Invitrogen) were applied for two hours at room temperature. Slides were then again washed three times with 1x PBS before coverslipping with Fluoromount G (0100-01, Southern Biotech) for analysis.

For EdU labeling, slides with optic nerve sections were incubated at room temperature for 30 minutes protected from light in freshly-prepared Alexa-647 EdU Cell Proliferation Assay (C10340, Thermo Fisher Scientific) cocktail after immunohistochemistry. Slides were washed three times in 1x PBS and coverslipped in Fluoromount-G (01001-01, Southern Biotech). For BODIPY 493/503 (D3922, Invitrogen) staining, slides were incubated in BODIPY for 15 minutes in 0.5µg/mL solution following application of secondary antibodies and then washed 3x in 1x PBS prior to coverslipping.

Retinal flatmounts were blocked in 10% fetal calf serum with 0.2% Triton-X100 for 1 hour with agitation. Retinae were then incubated with primary antibodies at 4°C with agitation in 1x PBS 0.2% Triton-X100. Primary antibodies included guinea pig anti-RBPMS (1:500; 1832, Phosphosolutions), rabbit anti-RBPMS (1:500 ABN1362, Millipore), mouse anti-β3 Tubulin (1:1000; T8660, Sigma), rabbit anti-phosphorylated c-Jun (1:500, 9261 Cell Signaling), mouse anti AP-2α (1:500, sc-12726 Santa Cruz), mouse anti-AP-2β (1:25, PCRP-TFAP2B-2E1, DSHB) and rabbit anti-cleaved caspase-3. Appropriate secondary antibodies were applied overnight with agitation at 4°C. To prepare for mounting on slides, each retina was cut with 4-5 incisions along the radial axis from the edge to about 2/3rds the distance to the optic disk then mounted on slides. Prolong Glass Antifade Mountant (P36980, Thermo Fisher Scientific) was applied prior to coverslipping.

### In situ hybridization

RNAscope *in situ* hybridization was used to detect *Rbpms* (527231, ACDBio) in combination with *Ecel1* (1137301-C2, ACDBio) or *Hrk* (475331-C3, ACDBio). The assay was performed according to the manufacturer’s instructions (RNAscope Multiplex Fluorescent V2 Assay, ACDBio). Briefly, 20 µm thick sections were mounted on Superfrost slides (1255015, Fisher Scientific) and stored at -80°C until *in situ* hybridization was performed. Slides were dehydrated in 50%, 70%, and 100% ethanol for five minutes before being placed into boiling target retrieval buffer for five minutes to unmask the target RNA. Slides were treated with H_2_O_2_ for 10 minutes at room temperature then Protease III was applied for 30 minutes at 40°C. Probes were hybridized for two hours at 40°C. Either *Ecel 1* or *Hrk* was assigned to channel 2 or 3 and diluted 1:50 in *Rbpms* probes assigned to channel 1. To test RNA integrity within the tissue, probes against housekeeping genes *Polr2a*, *Ppib* and *Ubc* (320881, ACDBio) were applied to slides cut in parallel along with 3-Plex negative control probes on an additional slide (320871, ACDBio). Signal amplification was performed according to the instructions of the kit. Signal detection utilized Opal520 (OP-001001, Akoya) and Opal 690 (OP-00106, Akoya), which were diluted 1:1000 in TSA buffer (322809, ACDBio). Nuclei were detected by DAPI stain applied for five minutes prior to coverslipping with Prolong Gold (P36934, Thermo Fisher Scientific).

### Electron microscopy tissue processing and analysis

Following postfixation, optic nerves were stored in a buffer of 1.5% paraformaldehyde, 1.5% glutaraldehyde, 50mM sucrose, 22.5mM CaCl_2_ 2H_2_O in 0.1M cacodylate buffer for at least seven days before embedding. Optic nerves were trimmed to two millimeters from the optic chiasm prior to plastic embedding and sections obtained approximately 50µm chiasmal of that to avoid dissection artefact. Optic nerves were post-fixed in 2% osmium tetroxide (19190, Electron Microscopy Sciences) with 1.5% potassium ferrocyanide (25154-20, Electron Microscopy Sciences) using a Biowave Pro+ microwave (Ted Pella). Contrast was enhanced by *en bloc* staining with 0.5% uranyl acetate (22400, Electron Microscopy Sciences) before dehydration in ethanol and embedding in Embed 812 (14120, Electron Microscopy Sciences). 0.5 µm sections were cut on an ultramicrotome and stained with 0.5% Toluidine Blue (22050, Electron Microscopy Sciences) with 0.5% sodium borate (21130, Electron Microscopy Sciences) to visualize the optic nerve for area measurements. 60nm sections were mounted on copper grids (T400-Cu, Electron Microscopy Sciences) then counter stained with 5% Uranyl Acetate for twenty minutes followed by Reynold’s Lead Citrate (80 mM Pb(NO₃)₂ 17900-25, Electron Microscopy Sciences and 120mM Sodium Citrate, 21140, Electron Microscopy Sciences) for six minutes. Grids were imaged at 4800x on a FEI Tecnai T12 transmission electron microscope with a 16 Mpx camera (Advanced Microscopy Techniques Corp). The density of axons and those with organelle accumulations was measured within ten random images per nerve. Organelle accumulations were counted if more than half the area of the axon was occupied by two or more organelles. We then multiplied the average density of axons with organelle accumulations within the optic nerve by the area measured on the adjacent Toluidine Blue section to get the total number of axons per nerve. For g-ratio analyses, 5-6 images were analyzed per animal at 4800x magnification. For quantifications at 10 and 20 weeks post tamoxifen, the axon and myelin were manually traced with the spline contour tool using the Zen 3.0 software (Zeiss) to determine the axon diameter relative to the axon diameter with myelin. For g-ratio measurements at eight weeks post tamoxifen, Myeltracer tool was used to calculate axon diameter relative to myelin diameter^73^. Demyelinated axons were not included in g-ratio analyses and analyses was conducted blinded to genotype.

### Serum neurofilament light chain (NfL) detection

After deep anesthesia with ketamine and xylazine as above, 0.5mL blood was acquired from mice immediately prior to perfusion via intracardiac puncture. Blood was allowed to clot for one hour before spinning at 500g for 10 minutes. The serum was removed and snap frozen on dry ice and stored at -80°C. NfL concentration was measured on the Simoa platform using the NF-light advantage kit V2 (Quanterix). To account for the high concentrations found in demyelinating mice that might go beyond the highest point of the calibrator, serum was bench-diluted to 1:4 or 1:8 (depending on available sample volume). On the Simoa, another 1:4 online dilution followed as part of the standard assay procedure, and final concentration was corrected for the applied dilution factor.

### Immunofluorescence image analysis

Immunostained sections were captured with a Zeiss ApoTome2 at 20x using 0.8NA lens. Optic nerve cross sections were imaged in their entirety with at least four sections 200 µm apart analyzed per mouse, per analysis. For cellular counts of OL lineage cells, the optic nerve was manually outlined using the spline contour tool in Zen 3.0 (Zeiss) and OLIG2-positive nuclei were counted first. Each OLIG2-positive cell was examined to see if it expresses CC1, PDGFRα, SYT4, NKX2.2 or EdU. For microglial and T-cell counts, only Iba1-positive or CD3-positive cells with DAPI nuclear staining were considered to be positive. For analysis of BCAS1, MBP, E06 and CD68 images were manually thresholded by an observer blinded to genotype and timepoint in Fiji ImageJ 1.53 (NIH) to determine the area occupied relative to the size of the optic nerve. For analysis of BODIPY immunofluorescence in microglia, a microglia/macrophage were first identified by IBA1+ staining then BODIPY fluorescence was measured within IBA1+ cells using ImageJ. Likewise, for E06, an IBA1+ mask was placed over the image and the area of E06 staining outside and inside microglia were quantified. To quantify phosphorylated c-Jun positive neurons in the spinal cord, whole spinal cords were imaged and the grey matter outlined using the NEUN channel and double positive cells were counted and examined to ensure there was DAPI present. At least four images of the lumbar spinal cord were quantified per animal.

Retinae were imaged in their entirety for analysis at 20x with 0.8NA lens using a Zeiss ApoTome2. To quantify RBPMS or phosphorylated c-Jun, cells were manually counted in the GCL within a 200 µm x 200 µm box placed at 500 µm, 1000 µm, 1500 µm and 2000 µm from the optic disk in each quadrant for a total of 16 regions. For quantifications of whether phosphorylated c-Jun+ cells were AP-2α/β or RBPMS-positive, each phosphorylated c-Jun positive cell was counted as above then determined if the cells were AP-2α/β or RBPMS positive. Cleaved caspase-3-positive cells were counted over the extent of the retina. For analysis of cleaved caspase-3 expression in AP2-α/β or RBPMS positive cells, each caspase-3 positive cell was individually evaluated for these markers per retina. The retina was manually outlined using the spline contour tool (Zen 3.0, Zeiss) to determine the area for density measurements. All analyses were conducted blind to genotype or treatment.

### Western blot

Mice were deeply anesthetized as above and perfused with 1x PBS before the optic nerve was removed and flash frozen on dry ice and stored at -80°C until protein was extracted. Thawed optic nerves were dounce homogenized in RIPA buffer along with complete protease inhibitors (11836153001, Roche) and phosphatase inhibitors (04906837001, Roche). Following homogenization, samples were spun at 13,000g for 15 minutes and the protein lysate was removed and frozen. Lysates from four nerves (two mice) were combined and protein was run on Bis-Tris-gel (NP0335BOX, Invitrogen). To transfer to a PVDF membrane (IPVH00010, Thermo Scientific), the gel blotting sandwich cassette (A25977, Thermo Fisher Scientific) was placed in transfer buffer (NP0006-01, Thermo Scientific) under 25V for 90 minutes. Following transfer, blots were rinsed in 1x TBS with 0.1% Tween-20 (TBST) before blocking in 1x TBST with 5% milk powder for one hour. Blots were probed with antibodies against DLK (GTX124127, Genetex), pMKK4 (9156, Cell Signaling), MKK4 (9152, Cell Signaling), pJNK (9251, Cell Signaling), JNK (9525, Cell Signaling), MOG (supernatant from clone 8-18C5, kind gift of R. Reynolds, Imperial College, London, UK), MBP (MAB386, Millipore) diluted in TBST with 2% BSA overnight (BP9706-100, Fisher Scientific). After overnight incubation, blots were washed in 1x TBST and incubated with appropriate HRP-conjugated secondary (Goat anti-rat 7077, Cell Signaling, Goat anti-mouse 7076, Cell Signaling, Goat anti-rabbit 7074, Cell Signaling) for two hours with 2% milk powder in TBST. Immunoreactivity was visualized using chemiluminescence (34080, Thermo Fisher Scientific) and imaged on a Syngene GBox iChemiXT. Blots were subsequently re-probed with β-actin-HRP as a loading control (A3854, Sigma). Densitometric analysis was performed in ImageJ 1.53 by quantifying the intensity of bands relative to loading control and then normalized relative to the mean of the Myrf^fl/fl^ control group.

### Laser capture microscopy

Eyes were dissected and snap frozen in OCT, then sectioned on a cryostat at 20 µm thickness and mounted onto Poly-L-Lysine (P1524, Sigma) coated membrane slides (414190-9041-000, Zeiss). Sections were fixed in 70% ethanol for two minutes before staining in Harris Modified Hematoxylin (HHS32, Sigma) with 0.2% glacial acetic acid for 30s. Sections were then immersed in 70% ethanol twice and 100% ethanol twice for 30s each and stored at -80°C in a sealed container until LCM was performed. A Zeiss Palm Microbeam microscope was used to conduct LCM with cut segments extracted onto the lid of adhesive cap tubes (415190-9181-000, Zeiss). The GCL was identified and sectioned—at least 20 sections per animal. For sampling of the whole retina, laser incisions were made through each retinal layer. Samples were treated with RLT lysis buffer and RNA isolated using the MicroRNAeasy kit (74004, Qiagen) as per the manufacturer’s instructions and frozen -80°C until sequencing.

### Bulk RNAseq

Following RNA isolation, RNA quantity and quality were evaluated on an Agilent 2100 Bioanalyzer using the Eukaryote Total RNA Pico. cDNA libraries were produced by loading 15ng of RNA for use with the Illumina Stranded Total RNA Prep, Ligation with Ribo-Zero Plus kit and sequenced with a NovaSeq 6000 at 50 million reads per sample. Raw reads were sorted based on barcodes and FASTQC files were produced. Reads were aligned to *Mus musculus* (GRCm38/Mm10) and expression counts were performed using STAR. DeSeq2 was run using Basepair software to determine differentially expressed genes between whole retina and the GCL or between the GCL of Myrf^ΔiSox10^ and Myrf^fl/fl^ mice. A total of six Myrf^fl/fl^ and six Myrf^ΔiSox10^ were compared for statistical analyses.

### Nuclei isolation and snRNAseq

Blood was removed from deeply anesthetized mice via intracardiac perfusion with 10mL of 1x PBS at 4-7 PM to reduce circadian fluctuations. Optic nerves were dissected out and immediately snap-frozen on dry ice. Frozen tissue was stored at -80°C for up to six months until subsequent processing. The nuclei isolation buffer (NIB, 146 mM NaCl, 5 mM Tris-HCl, 1 mM CaCl_2_, 21 mM MgCl_2_, 0.03% Tween-20, 0.01% BSA, 1 µg/mL actinomycin D, pH 7.5) was prepared with one tablet of protein inhibitor cocktail (cOMPLETE Mini lacking EDTA, 11873580001, Roche) along with 15uL of RNAsin (N2615, Promega) per 10 mL NIB. When optic nerves were removed from the freezer, they were immediately placed in cooled 2mL NIB solution in a 7mL Dounce grinder and ground 20 times with a loose pestle. Then, the homogenate was passed through a 200 µm strainer (43-50200-03, Pluriselect). The homogenate was ground 10 additional times with a tight pestle, then 2 mL NIB was added and the homogenate was passed through a 40 µm strainer (43-50040-03, Pluriselect). The homogenate was ground five times with a tight pestle (B), then passed through a 20 µm filter (43-50020-03, Pluriselect). The sample was then centrifuged at 500g for five minutes at 4°C three times with the supernatant discarded and new NIB added each time. Following the last centrifugation step, the pellet was resuspended in 0.5 mL NIB with 5 µL SuperaseIN (AM2696, Thermo Fisher Scientific) along with 1% BSA and mixed 1:200 with RedDot (40060, Biotium). The samples were then isolated from debris by fluorescence-activated nuclei sorting (FANS). Two main gates were used: a 638 emission for the RedDot stain and a low trigger pulse width as singlet discriminator (Supplementary Fig. 12). 100,000 nuclei were aimed to be sorted. Following sorting, samples were centrifuged at 300g for one minute at 4°C, held on ice for one minute then spun for one minute at 300g at 4°C. The top supernatant was carefully removed and the nuclei were then prepared for snRNAseq using a Chromium Next GEM Single Cell 3′ Reagent Kit v3.1 (10x Genomics). Single nuclei were partitioned in droplets with single gel beads, which contained primers with cell-tagging indexes. Single nucleus suspensions were targeted to 10,000 nuclei per sample with 500 million reads per sample. The resulting cDNA was used as a template for library preparation. Samples were sequenced using a NovaSeq 6000 and FASTQ files were prepared using bcl2fastq (Illumina) and then aligned to the mouse GRCm38/mm10 reference genome using Cellranger (v7.0.0, 10x Genomics). Reads were mapped to both exonic and intronic regions.

### snRNA-seq analyses

Data was analyzed using R (v4.2.1) and the Seurat package (v4.3.0)^74^. First, quality control of each sample was performed. Ambient RNA was removed using SoupX^75^ (v1.6.2). To remove doublets and debris, nuclei were filtered based on 1,000 < nFeature_RNA < 4,000 as well as 1,250 < nCount_RNA < 10,000. Mitochondrial genes were removed by manually excluding all features starting with “mt-“. One sample (a Myrf^fl/fl^ sample) failed quality control and was not included in further analysis. Following quality control, the seven samples were normalized with ‘*SCTransform^76^*’ (v0.3.5) and integrated using ‘*FindIntegrationAnchors*’ function in Seurat. The integrated dataset was visualized by UMAP plotting techniques. Nuclei were first clustered using 35 principal component dimensions. Differentially expressed genes (DEGs) were identified using Seurat’s ‘*FindMarkers*’ with the following analysis criteria: Log2 fold change > 0.322, the minimum percentage of nuclei expressing the gene = 0.25, and adjusted p-value (calculated with the Wilcox significance test) < 0.05. Top genes were used to identify each cluster.

Several samples had neuronal nuclear contamination from the dissection indicated by nuclei with high *Rbfox3, Syt7*, and *Snap25* and these clusters were manually excluded. After the neuronal contamination was removed, we re-clustered using 33 principal component dimensions to account for the removal of a highly diverse neuronal population that may have affected the original UMAP clustering. We re-ran the DEG analysis for the final UMAP clustering using the analysis criteria stated above and annotated the clusters using established markers. Next, we compared the DEGs between the control mice derived from different lines (Myrf^fl/fl^; Plp1-CreERT-negative and Myrf^fl/fl^; Sox10CreERT-negative) using the analysis criteria stated. Because both control lines had less than 9 DEGs in total, for the ease of analysis between genotypes, we combined the three control samples (hereby referred to as Myrf^fl/fl^). After combining the controls, DEGs between Myrf^fl/fl^ and Myrf^ΔiPlp1^ or Myrf^ΔiSox10^ were identified with the analysis criteria. Likewise, DEGs between the two knock-out lines (Myrf^ΔiPlp1^ or Myrf^ΔiSox10^) were identified using the analysis criteria above. DEGs for KOOLs were then displayed using a volcano plot (ggplot2 3.42). and transcripts with log2 fold change > 0.50 were highlighted as enriched in Myrf^ΔiPlp1^ or Myrf^ΔiSox10^ nuclei.

For microglia re-clustering, we subdivided these cells and reclustered using eight principal component dimensions based off the elbowplot. DEGs for microglia subclusters were identified with the same analysis criteria as above. DIM3 population DEGs were calculated with the following analysis criteria: log2 fold change > 0.50 and visualized with a volcano plot (ggplot2 3.42). Gene set enrichment analysis (GSEA) for molecular function pathways was performed on the DAM3 cluster differentially expressed genes (p adjusted < 0.05) using ClusterProfiler (v4.6.2). Trajectory pathway analysis in psuedotime was performed on the OL lineage using Monocle3 (v1.3.1).

### Production of viral constructs

SgRNAs were designed using CRISPROn^77^ and compared to previously published sgRNAs^78^. Five sgRNAs for *Map3K12* and *Map3K13* were tested in total for indel formation in cultured Neuro2a cells expressing Cas9. Px333 (Addgene # 64073) was modified by restriction enzymes to insert oligonucleotides with the MluI site and ApaI site at the XhoI and KpnI sites, respectively. This allows the removal of tandem U6 promoters with sgRNAs and insertion into AAV-U6-sgRNA-hSyn-mCherry (Addgene #87916) in place of the single U6-sgRNA when digested with MluI and ApaI. SgRNAs were tested in the modified px333 plasmid and validated by TIDEs (tracking of indels by decomposition)^79^. The two best sgRNAs were chosen based on the degree of decomposition after the PAM site. While all sgRNAs tested demonstrated decomposition after the PAM site, the sgRNAs which were used in the study to target *Map3K12* at exons 4 and 9 had the highest decomposition and have the following sequences AGGGTGTTCGGGTTTCATGG and TGTAGAGAGCACATCAGCGG. The guides utilized against *Map3K13* had the sequences TCTGGGGAACAGCAACACTG and GGTCACGGTGTATAATCTTG^78^ targeting exons 3 and 5 of *Map3K13*, respectively. SgRNAs against LacZ and GFP for the control AAV were taken from previously validated sgRNAs^80, 81^. AAV2-hSyn1-Cre was generated by replacing the mCherry open reading frame from AAV-U6-sgRNA-hSyn-mCherry (Addgene #87916) with the Cre open reading frame from AAV:ITR-U6-sgRNA-pCBh-Cre-WPRE-hGHpA-ITR (Addgene #60229). Large-scale packaging into AAV2 of viral vectors for intravitreal injection was completed by Vector Biolabs (AAVs with sgRNAs against *Map3K12*, *Map3K13* or *Map3K12*/*Map3K13*) or Vector Biosystems (sgLacZ/sgGFP) or using the OHSU Molecular Virology Core (AAV2-hSyn1-Cre).

### Cas9-expressing Neuro2a cell culture

Mouse Neuro2a cells (CCL-131, ATCC) expressing Cas9 (cells were transduced with Cas9 in the pQXCIH plasmid) were grown in Dulbeco’s modified Eagle medium (11960-044, Gibco) (DMEM) with 10% FBS supplemented with glutamine (25030-081, Gibco), Penicillin-Streptomycin (100 U/mL penicillin, 100 mg/mL streptomycin;15140-122, Gibco); and Sodium Pyruvate (11360-070, Gibco). Neuro2a cells were incubated at 37 °C and 5% CO_2_ and were passaged every three days. To transfect Neuro2a cells with sgRNA expressing plasmids, we used Lipofectamine 2000 (52758, Invitrogen) and the cells were collected 24 hours later and DNA extracted using DNeasy Blood and Tissue Kit as per the manufacturer’s instructions (69504, Qiagen).

### Statistics

Statistical analyses were conducted with Prism 10.2.2 (Graphpad) or SPSS v13.0 and all data is presented as mean ± standard error. Sample sizes are indicated by the number of dots in the figures or are otherwise explicitly stated. All tests are two-sided. To test for normality, the Shapiro-Wilk test was used. To test for homogeneity of variance, Levene’s test was used for comparison of two groups and Brown-Forsythe test for three or more. If data was not normally distributed appropriate non-parametric tests were run. If a time point had less groups than others in the analysis because of premature euthanasia of Myrf^ΔiSox10^ mice, this time point was treated as an independent experiment for statistics. For all statistical analyses *n* represents a single animal, except for the optic nerve snRNAseq and western blots, where two animals worth of nerves were combined per sample (n). Degree of significance was indicated in figures by * = p < 0.05, ** p < 0.01, *** p< 0.001 or **** p < 0.0001 unless stated otherwise. Animals were assigned to group based on genotype or to treatment by random selection. The number of mice used in experiments was based on previous publications with similar methodology. Statistical comparisons are outlined in table S6 and *t*, *F, H* statistics, *P* values and degrees of freedom reported when appropriate.

## Materials availability

Plasmids generated in this study for DLK or LZK knockout have been deposited at Addgene (Plasmid #208834, 208835, 208836, 208837) and AAV2-hSyn1-Cre is available upon reasonable request. Transgenic mouse lines are available at Jackson laboratory (Sox10-CreERT JAX #027651, Plp1-CreERT JAX #005975, Myrf^fl/fl^ JAX #010607, Rosa26-Cas9 mice JAX #024858, Sun1-GFP mice JAX #030952) and the Myrf^fl/fl^ line on a C57BL/6 background is available from the Emery lab upon reasonable request. All other reagents were from commercial sources.

## Data availability

SnRNA-seq data has been deposited in the NCBI Gene Expression Omnibus (GEO; http://www.ncbi.nlm.nih.gov/geo/) and is accessible through GEO Series accession number GSE243788. SnRNA-Seq data can be viewed in our interactive browser at https://emerylab.shinyapps.io/Myrf_iCKO_OpticNerve/. Bulk RNA-Seq data has been deposited in the NCBI GEO and is accessible through GEO Series accession number GSE245362. Original data are available from the authors upon reasonable request.

## Code availability

All original code has been deposited at (Github: https://github.com/EmeryLab/Myrf_iCKO_OpticNerve).

