## Supplementary Figures 1-12 for "Remyelination protects neurons from DLK-mediated neurodegeneration"

Supplementary materials for '*Remyelination protects neurons from DLK-mediated neurodegeneration*.'

Contents include the following:

Supplementary Figs 1-12

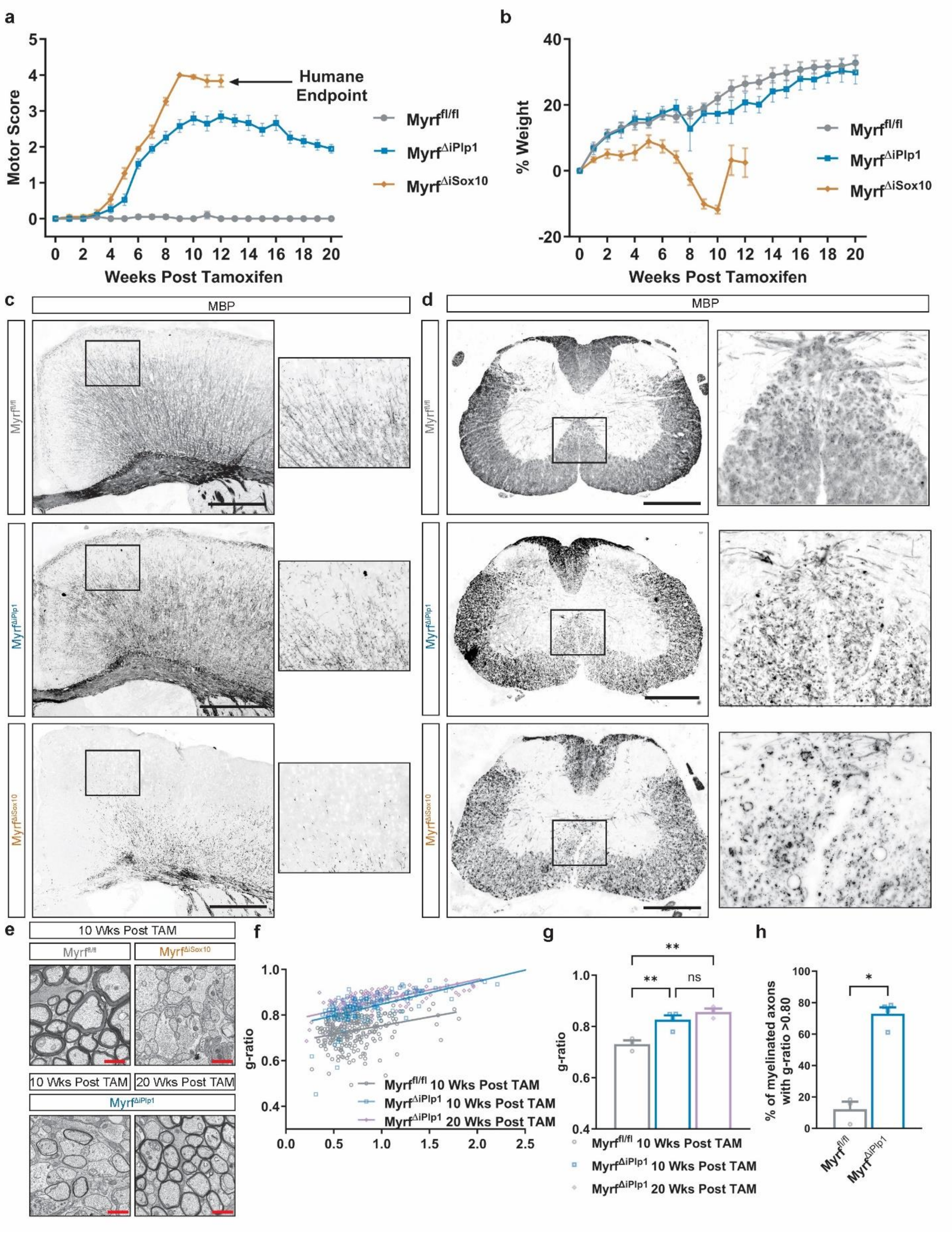
a
Motor Score
Weeks Post Tamoxifen
Humane Endpoint
Myrffl/fl
Myrf∆iPlp1
Myrf∆iSox10
b
% Weight
Weeks Post Tamoxifen
Myrffl/fl
Myrf∆iPlp1
Myrf∆iSox10
c
MBP
Myrffl/fl
Myrf∆iPlp1
Myrf∆iSox10
d
MBP
Myrffl/fl
Myrf∆iPlp1
Myrf∆iSox10
e
10 Wks Post TAM
Myrffl/fl
Myrf∆iSox10
10 Wks Post TAM
20 Wks Post TAM
Myrf∆iPlp1
f
g-ratio
Myrffl/fl 10 Wks Post TAM
Myrf∆iPlp1 10 Wks Post TAM
Myrf∆iPlp1 20 Wks Post TAM
g
g-ratio
ns
Myrffl/fl 10 Wks Post TAM
Myrf∆iPlp1 10 Wks Post TAM
Myrf∆iPlp1 20 Wks Post TAM
h
% of myelinated axons with g-ratio >0.80
Myrffl/fl
Myrf∆iPlp1

**Supplementary Fig.1: Motor behavioral deficits are associated with CNS-wide demyelination and recovery is associated with remyelination in Myrf$^{\Delta iPlp1}$ mice.**

**a** Motor scores. 0 = no deficit, 1 = limp tail in gait, 2 = hindlimb weakness, 3 = ataxia and severe hindlimb weakness, 4 = hindlimb paresis, 5 = hindlimb paralysis. Myrf$^{\Delta iSox10}$ mice have worsened motor scores relative to Myrf$^{\Delta iPlp1}$ mice ($P = 0.0039$) and both Myrf$^{\Delta iPlp1}$ and Myrf$^{\Delta iSox10}$ mice have worsened motor scores relative to Myrf$^{fl/fl}$ ($P < 0.0001$). **b** Quantification of percent weight loss in each strain. Myrf$^{\Delta iSox10}$ mice have increased weight loss relative to Myrf$^{fl/fl}$ and Myrf$^{\Delta iPlp1}$ ($P < 0.0001$ for both comparisons). **c** MBP in the cortex and corpus callosum at 10 weeks post tamoxifen. The medial corpus callosum and upper layer of the cortex are prominently demyelinated in Myrf$^{\Delta iSox10}$ mice, and the upper layers of the cortex feature a notably lower density of myelin in Myrf$^{\Delta iPlp1}$ mice relative to Myrf$^{fl/fl}$. Boxed regions are of the upper cortical layers. **d** Lumbar spinal cord with MBP immunohistochemistry. There is degenerating myelin and demyelination at 10 weeks post tamoxifen in both Myrf$^{\Delta iPlp1}$ and Myrf$^{\Delta iSox10}$ mice. Boxed region is the ventral white matter in Myrf$^{\Delta iPlp1}$ and Myrf$^{\Delta iSox10}$ mice. **e** Example high-magnification electron micrographs of axons within the optic nerve of Myrf$^{fl/fl}$, Myrf$^{\Delta iPlp1}$ and Myrf$^{\Delta iSox10}$ mice. **f** g-ratio of myelinated axons in Myrf$^{fl/fl}$ and Myrf$^{\Delta iPlp1}$ mice. **g** Average g-ratio of myelinated axons indicates axons in Myrf$^{\Delta iPlp1}$ mice are thinly myelinated relative to Myrf$^{fl/fl}$ at both 10 ($p = 0.0070$) and 20 ($p = 0.0023$) weeks following tamoxifen administration. Axons lacking myelin were excluded from g-ratio analysis. 552 axons were analyzed. **h** Percentage of myelinated axons with a g-ratio > 0.80 in Myrf$^{fl/fl}$ and Myrf$^{\Delta iPlp1}$ at 10 weeks post tamoxifen. Axons with a g-ratio of > 0.80 are rare in Myrf$^{fl/fl}$ relative to Myrf$^{\Delta iPlp1}$ ($P = 0.0286$).Repeated measures mixed effect model with Tukey's *post hoc* test in **a**, **b**. One-way ANOVA with Tukey's *post hoc* test in **g** and Mann-Whitney U in **h**. Scale bars are 500 µm in **c** and **d** and 1 µm in **a**. Error bars are SEM.

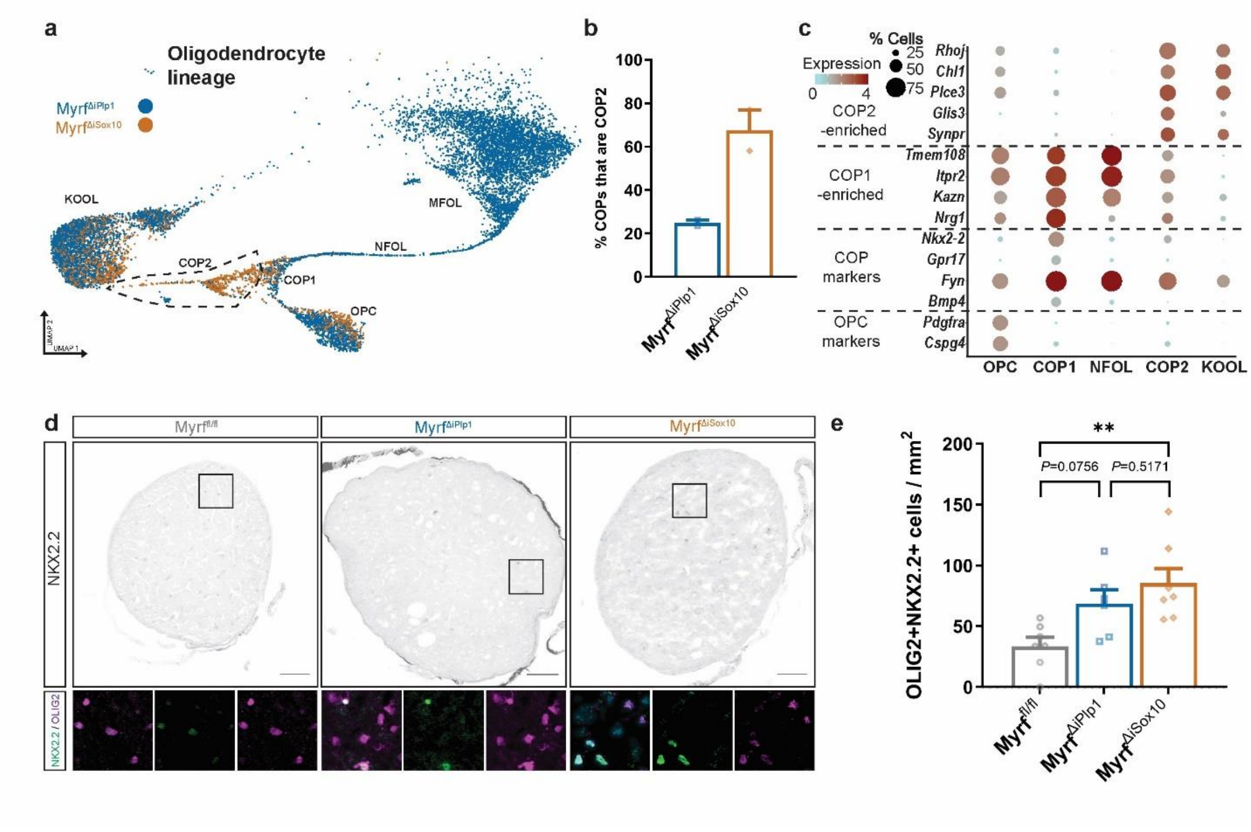


**Supplementary Fig. 2: Transcriptionally distinct COPs are formed in Myrf$^{\Delta iSox10}$ mice.**

**a** UMAP of the OL lineage in Myrf$^{\Delta iPlp1}$ and Myrf$^{\Delta iSox10}$ mice. Myrf$^{\Delta iSox10}$ mice show an enrichment of the COP2 cluster. **b** Ratio of COPs that are COP2 in Myrf$^{\Delta iPlp1}$ and Myrf$^{\Delta iSox10}$ optic nerves. **c** Dot plot of OPC, COP1, COP2, NFOL and KOOL clusters for transcripts enriched in OPC, COP1 and COP2. COP1-enriched transcripts are often co-expressed by NFOLs whereas COP2-enriched transcripts are often co-expressed by KOOLs. **d** Optic nerves stained with NKX2.2, which is enriched in COP1 and COP2. Boxed areas are shown below along with OLIG2, a pan-OL lineage marker. **e** OLIG2+ and strongly-expressing NKX2.2+ cells are enriched in Myrf$^{\Delta iSox10}$ relative to Myrf$^{fl/fl}$ ($P = 0.0058$). One-way ANOVA with Tukey's *post hoc* test. Error bars are SEM. Scale bars are 50 µm in **d**.

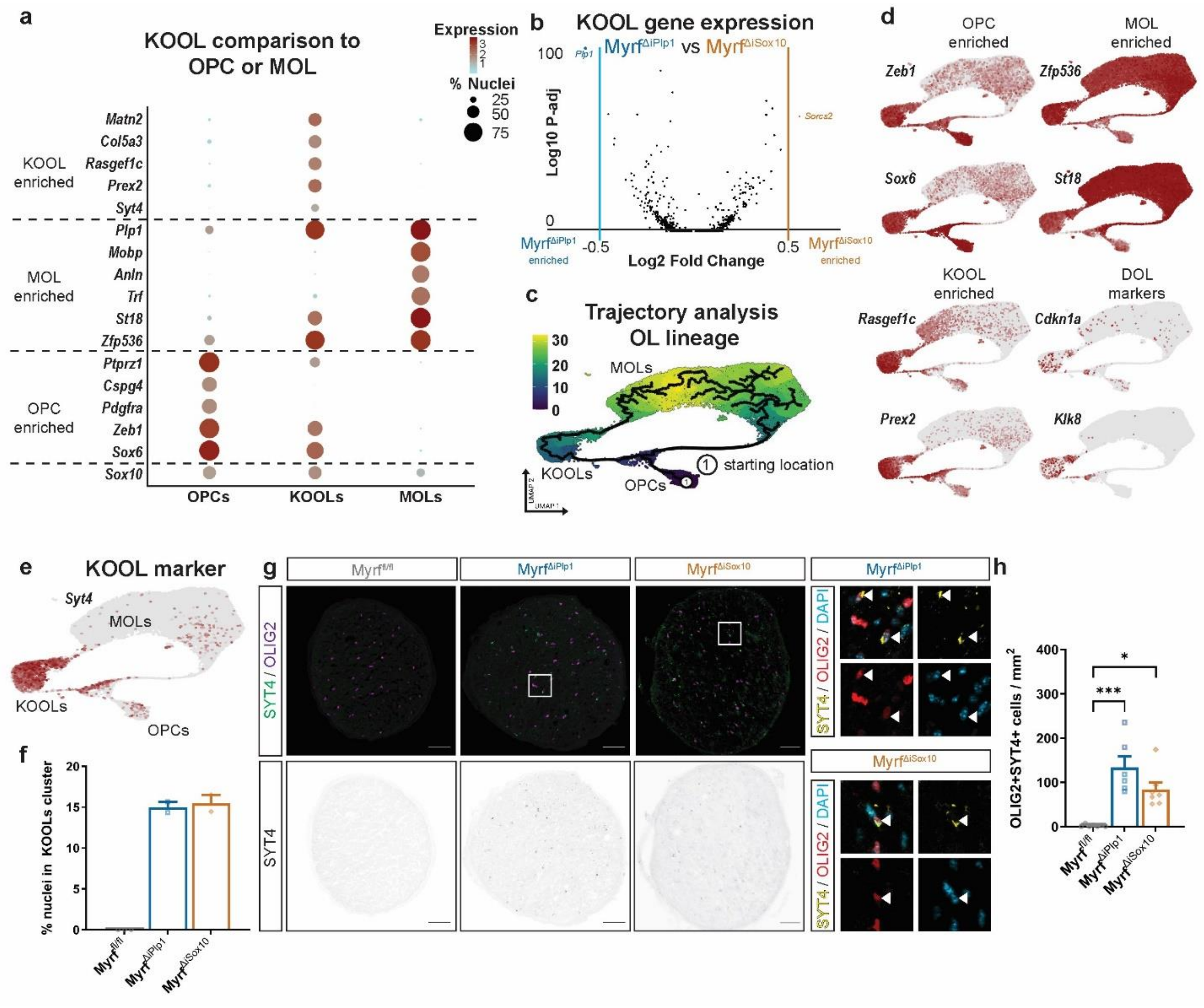
a
KOOL comparison to OPC or MOL
Expression
% Nuclei
25
50
75
KOOL enriched
Matn2
Col5a3
Rasgef1c
Prex2
Syt4
MOL enriched
Plp1
Mobp
Anln
Trf
St18
Zfp536
OPC enriched
Ptprz1
Cspg4
Pdgfra
Zeb1
Sox6
Sox10
OPCs
KOOLs
MOLs
b
KOOL gene expression
Myrf^ΔiPlp1 vs Myrf^ΔiSox10
Log10 P-adj
Log2 Fold Change
Plp1
Sorcs2
Myrf^ΔiPlp1 enriched
Myrf^ΔiSox10 enriched
c
Trajectory analysis OL lineage
MOLs
KOOLs
OPCs
starting location
d
OPC enriched
MOL enriched
KOOL enriched
DOL markers
Zeb1
Zfp536
Sox6
St18
Rasgef1c
Cdkn1a
Prex2
Klk8
e
KOOL marker
Syt4
f
% nuclei in KOOLs cluster
Myrf^fl/fl
Myrf^ΔiPlp1
Myrf^ΔiSox10
g
SYT4 / OLIG2
SYT4
SYT4 / OLIG2 / DAPI
h
OLIG2+SYT4+ cells / mm²

**Supplementary Fig. 3: KOOLs are a transcriptionally unique population of OL lineage cells selective to *Myrf* knockout mice.**

**a** Dot plot of comparison of key transcripts enriched in KOOL, OPC or MOL clusters. **b** Volcano plot of transcripts differentially expressed between Myrf$^{\Delta iSox10}$ and Myrf$^{\Delta iPlp1}$ KOOLs. $Log_2$(fold change) > 0.5 adjusted $p<0.05$. Wilcoxon rank-sum test. Just two transcripts meet this criteria (*Plp1* and *Sorcs2*). **c** Trajectory analysis of OL lineage starting at OPCs. **d** UMAPs for individual transcripts expressed in KOOLs and enriched in OPCs, OLs, KOOLs, or markers upregulated during cellular stress (DOLs: disease-associated OLs). **e** UMAP of *Syt4*, demonstrating enriched expression in KOOLs. **f** Percentage of nuclei in KOOL cluster from snRNAseq of the optic nerve. Both Myrf$^{\Delta iSox10}$ and Myrf$^{\Delta iPlp1}$ have a population of nuclei clustered as KOOLs whereas they are nearly absent in Myrf$^{fl/fl}$. **g** SYT4 and OLIG2 staining in the optic nerve at ten weeks post tamoxifen. Boxed areas in Myrf$^{\Delta iPlp1}$ and Myrf$^{\Delta iSox10}$ are enlarged to the right. SYT4 is closely-associated with OLIG2+ cells. **h** OLIG2+ SYT4+ cells are enriched in the optic nerves of both Myrf$^{\Delta iPlp1}$ ($P = 0.0004$) and Myrf$^{\Delta iSox10}$ ($P = 0.0379$) mice at 10 weeks post tamoxifen. Kruskal-Wallis test with Dunn's test to compare individual groups. Error bars are SEM. Scale bars are 50 µm in **g**.

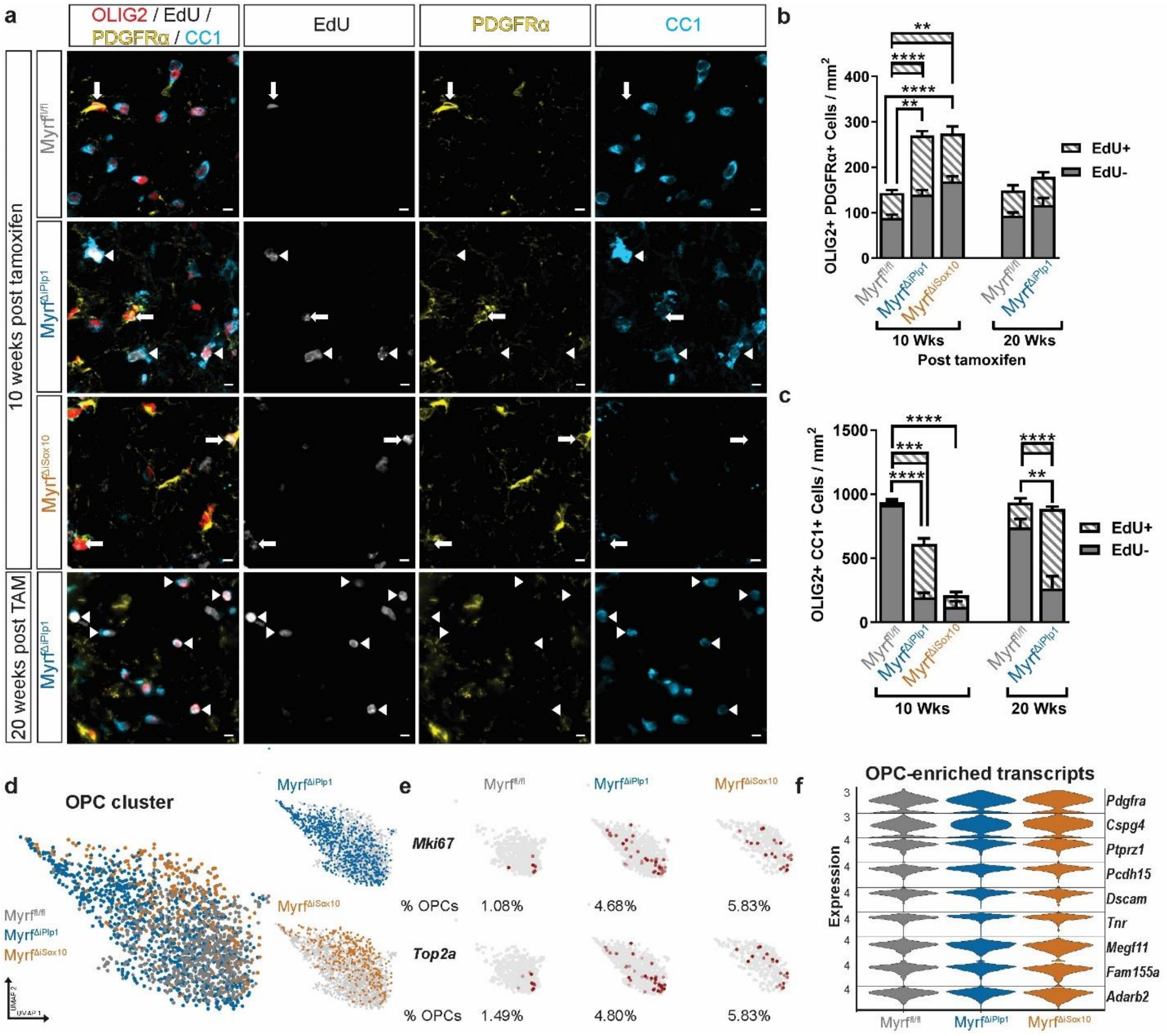
a
OLIG2 / EdU / PDGFRα / CC1
EdU
PDGFRα
CC1
10 weeks post tamoxifen
20 weeks post TAM
Myrf^fl/fl
Myrf^ΔiPlp1
Myrf^ΔiSox10
b
OLIG2+ PDGFRα+ Cells / mm²
EdU+
EdU-
10 Wks
20 Wks
Post tamoxifen
c
OLIG2+ CC1+ Cells / mm²
d
OPC cluster
e
Mki67
% OPCs 1.08% 4.68% 5.83%
Top2a
% OPCs 1.49% 4.80% 5.83%
f
OPC-enriched transcripts
Expression
Pdgfra
Cspg4
Ptprz1
Pcdh15
Dscam
Tnr
Megf11
Fam155a
Adarb2

**Supplementary Fig. 4: OPC proliferation does not differ between Myrf$^{\Delta iPlp1}$ and Myrf$^{\Delta iSox10}$ mice.**

**a** High magnification images of optic nerves stained for OLIG2, CC1, PDGFRα and EdU. Arrows indicate EdU+PDGFRα+ cells and arrowheads indicate EdU+CC1+ cells. **b** Quantification of OLIG2 PDGFRα double-positive OPCs with and without EdU. Striated significance bars indicate comparison between EdU+ cells, and unstriated bars between EdU-negative cells. There is an increase in the density of OLIG2+PDGFRα+ EdU- cells at 10 weeks post tamoxifen in Myrf$^{\Delta iPlp1}$ (*P* = 0.0029) and Myrf$^{\Delta iSox10}$ (*P* < 0.0001) mice relative to Myrf$^{fl/fl}$ mice. There is also an increase in the density of OLIG2+PDGFRα+ EdU+ cells in Myrf$^{\Delta iPlp1}$ mice (*P* < 0.0001) and Myrf$^{\Delta iSox10}$ (*P* = 0.0031) relative to Myrf$^{fl/fl}$ but Myrf$^{\Delta iSox10}$ and Myrf$^{\Delta iPlp}$ do not differ (*P* = 0.2895). **c** Density of OLIG2+CC1+ cells with and without EdU. Incorporation of EdU into OLIG2+CC1+ cells is indicative of oligodendrogenesis. There is a decrease in the number of OLIG2+CC1+EdU- cells in Myrf$^{\Delta iPlp1}$ and Myrf$^{\Delta iSox10}$ relative to Myrf$^{fl/fl}$ (*P* < 0.0001) but no difference between Myrf$^{\Delta iPlp1}$ and Myrf$^{\Delta iSox10}$ (*P* = 0.6623). There is an increase in OLIG2+CC1+EdU+ cells in Myrf$^{\Delta iPlp1}$ relative to Myrf$^{fl/fl}$ at 10 weeks (*P* = 0.0003) and 20 weeks post tamoxifen (*P* < 0.0001). **d** UMAP of OPCs split by genotype. **e** OPC cluster separated by genotype with transcript expression of proliferation markers *Mki67* (transcript for KI67 protein) or *Top2a* visualized. **f** Violin plot of transcripts highly enriched in OPCs split by genotype. There are broadly similar levels of expression of these transcripts between genotypes in OPCs. For EdU-negative cells in **b** and **c**, a one-way ANOVA with Tukey's *post hoc* for individual comparisons was run and to compare groups at 10 weeks post tamoxifen and a Student's t-test was run to compare groups at 20 weeks post tamoxifen in EdU negative and positive cells. At ten weeks post TAM in EdU-positive cells, a one-way ANOVA with Tukey's *post hoc* was used for pairwise comparisons in **b** and a Kruskal Wallis with Dunn's test was used in **c**. Error bars are SEM. Scale bar is 5 µm in **a**.

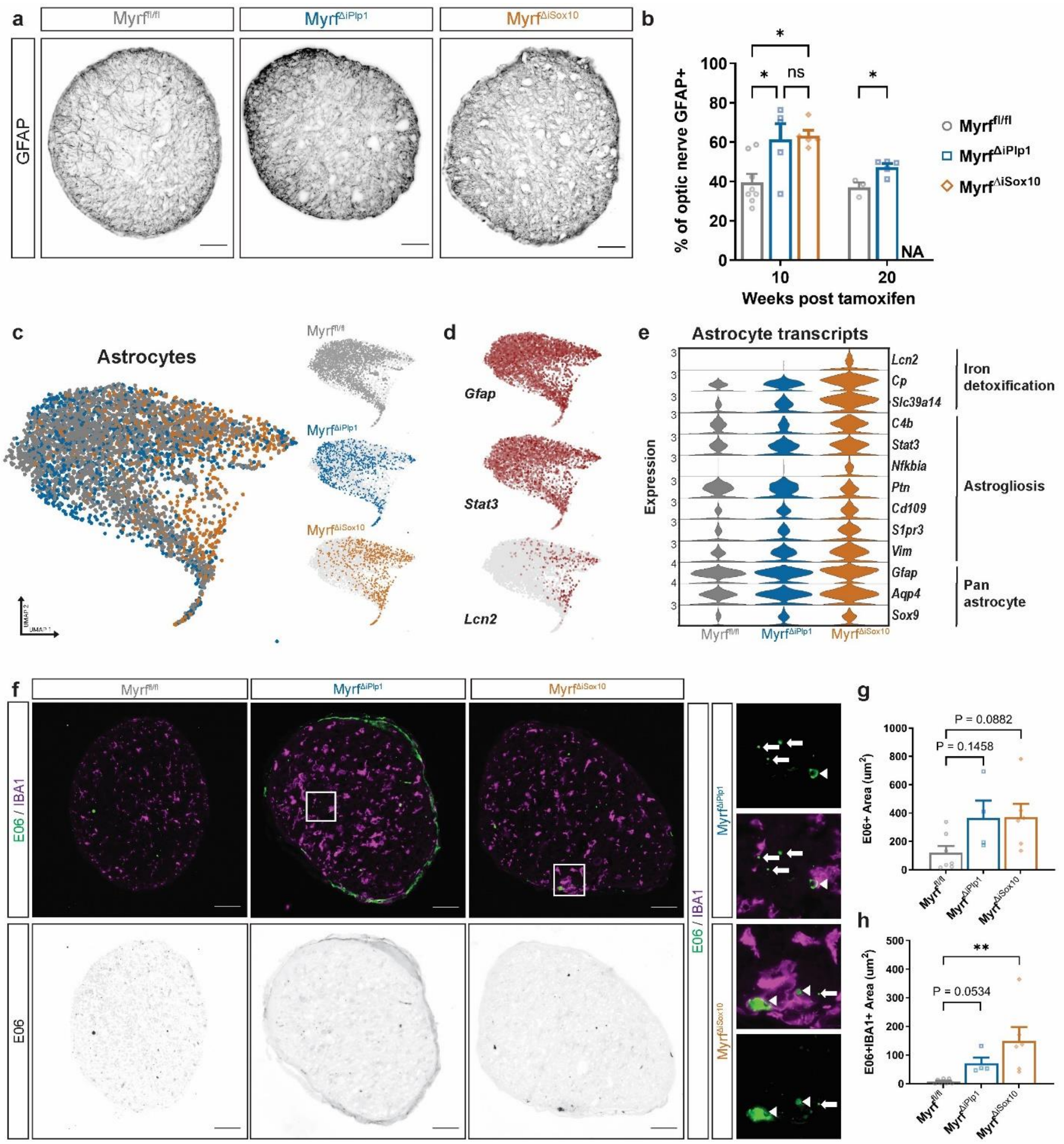
a
Myrf fl/fl
Myrf ΔiPlp1
Myrf ΔiSox10
GFAP
b
% of optic nerve GFAP+
Weeks post tamoxifen
ns
NA
Myrf fl/fl
Myrf ΔiPlp1
Myrf ΔiSox10
c
Astrocytes
d
Gfap
Stat3
Lcn2
e
Astrocyte transcripts
Expression
Lcn2
Cp
Slc39a14
C4b
Stat3
Nfkbia
Ptn
Cd109
S1pr3
Vim
Gfap
Aqp4
Sox9
Iron detoxification
Astrogliosis
Pan astrocyte
f
E06 / IBA1
E06
g
E06+ Area (um2)
P = 0.0882
P = 0.1458
h
E06+IBA1+ Area (um2)
P = 0.0534
**

**Supplementary Fig. 5: Astrogliosis and upregulation of iron detoxification transcripts following remyelination failure in Myrf$^{\Delta iSox10}$ mice.**

**a** Micrographs of GFAP staining in the optic nerve of Myrf$^{fl/fl}$, Myrf$^{\Delta iPlp1}$ and Myrf$^{\Delta iSox10}$ mice at 10 weeks post tamoxifen. **b** Quantification of the percentage of the optic nerve that is GFAP+. There is an increase in GFAP reactivity in Myrf$^{\Delta iPlp1}$ mice at 10 weeks post tamoxifen ($P$ = 0.0223) and 20 weeks post tamoxifen ($P$ = 0.0126) relative to Myrf$^{fl/fl}$. Myrf$^{\Delta iSox10}$ mice also have more GFAP reactivity relative to Myrf$^{fl/fl}$ ($P$ = 0.0133) but do not differ from Myrf$^{\Delta iPlp1}$ mice ($P$ = 0.9704). **c** UMAP of astrocytes broken down by genotype. **d** UMAPs of *Gfap, Stat3 and Lcn2* in astrocytes. *Lcn2* transcript expression overlaps with Myrf$^{\Delta iSox10}$ astrocytes. **e** Violin plot of known astrocyte markers and genes enriched during astrogliosis. Myrf$^{\Delta iSox10}$ astrocytes upregulate iron processing and detoxification genes including *Lcn2*, *Cp* and *Slc39a14*. **f** Optic nerves at 10 weeks post tamoxifen stained for IBA1+ microglia/macrophages and E06, a marker of oxidized phospholipids. Boxed areas shown in high magnification to the right with E06 staining within IBA1+ cells indicated by arrowheads or outside of IBA1+ cells (indicated by arrows) in Myrf$^{\Delta iPlp1}$ and Myrf$^{\Delta iSox10}$ mice. **g** Quantification of total E06 area in optic nerve demonstrates no statistically significant upregulation in Myrf$^{\Delta iPlp1}$ ($P$ = 0.1458) and Myrf$^{\Delta iSox10}$ ($P$ = 0.0882) mice at ten weeks post tamoxifen. **h** E06+ within IBA1+ cells is upregulated in Myrf$^{\Delta iSox10}$ relative to Myrf$^{fl/fl}$ ($P$ = 0.0033). One-way ANOVA with Tukey's *post hoc* used **g** and for comparisons at week 10 in **b**. Kruskal Wallis test with Dunn's test in **h**. Student's t-test for comparison at week 20 in **b**. ns= not statistically significant. Scale bar is 50 µm in **a** and **f**.

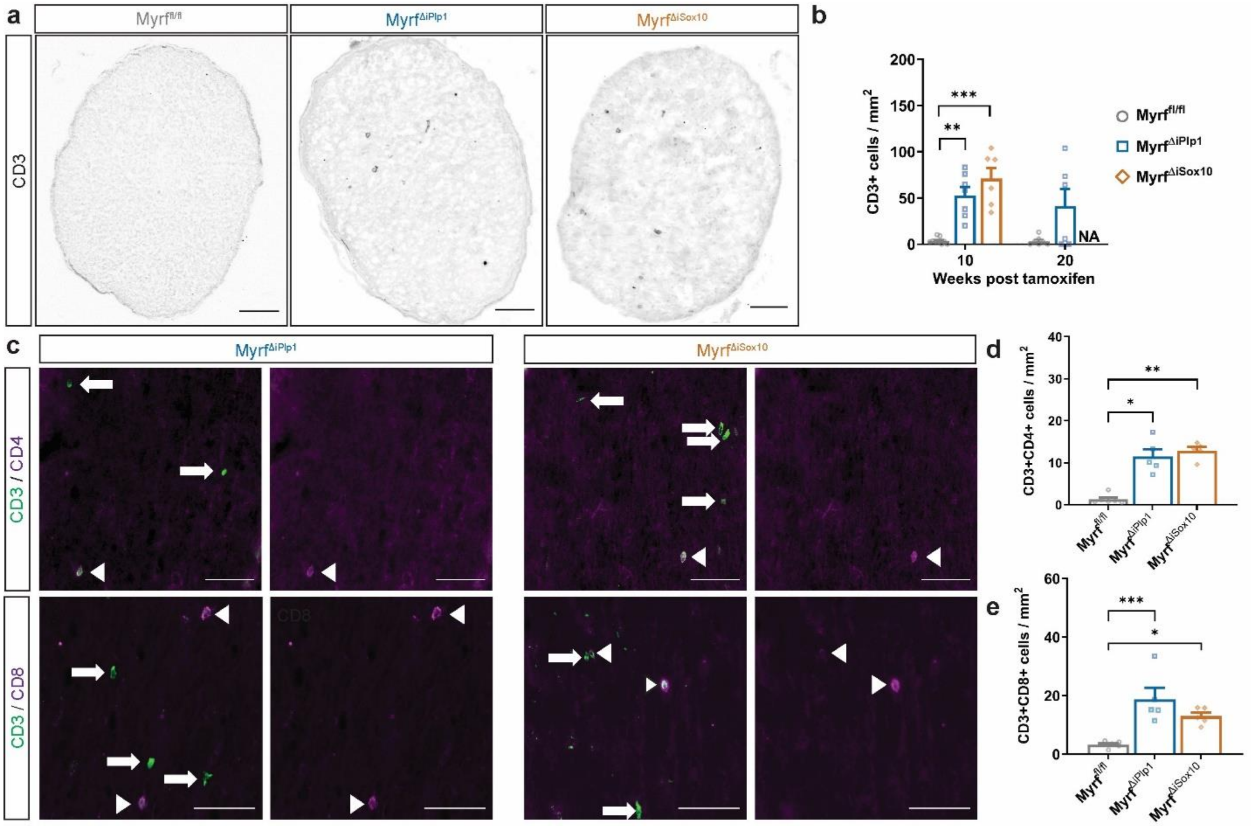


**Supplementary Fig 6: T cells infiltrate the optic nerve following demyelination in Myrf$^{\Delta iPlp1}$ and Myrf$^{\Delta iSox10}$ mice but do not differ in number or subtype.**

**a** Overview images of the optic nerves stained with CD3 demonstrates T cell infiltration into the parenchyma during demyelination in Myrf$^{\Delta iPlp1}$ and Myrf$^{\Delta iSox10}$ mice. **b** Quantification of CD3+ cells in the optic nerve. There is increased density of CD3+ cells at 10 weeks post tamoxifen in Myrf$^{\Delta iPlp1}$ ($P = 0.0090$) and Myrf$^{\Delta iSox10}$ mice ($P = 0.0008$) relative to Myrf$^{fl/fl}$ mice but Myrf$^{\Delta iPlp1}$ and Myrf$^{\Delta iSox10}$ mice do not differ from each other ($P > 0.9999$). **c** High magnification images of CD3, CD4 and CD8 positive T-cells in the optic nerve at 10 weeks post tamoxifen of Myrf$^{\Delta iPlp1}$ and Myrf$^{\Delta iSox10}$ mice. Arrowheads label CD4 or CD8+ cells and arrows label CD3+ cells without CD4 or CD8. **d** CD3+CD4+ cells increase in both Myrf$^{\Delta iPlp1}$ ($P = 0.0414$) and Myrf$^{\Delta iSox10}$ mice ($P = 0.0061$) relative to Myrf$^{fl/fl}$ but do not differ from each other ($P > 0.9999$). **e** CD3+CD8+ density is increased in both Myrf$^{\Delta iPlp1}$ ($P = 0.0006$) and Myrf$^{\Delta iSox10}$ mice ($P = 0.0201$) but do not differ from each other ($P = 0.2087$). Kruskal Wallis test with Dunn's test in **b** and **d.** One way ANOVA with Tukey's *post hoc* for individual comparisons in **e**. Error bars are SEM. Scale bar is 50 µm in **a** and **c**.

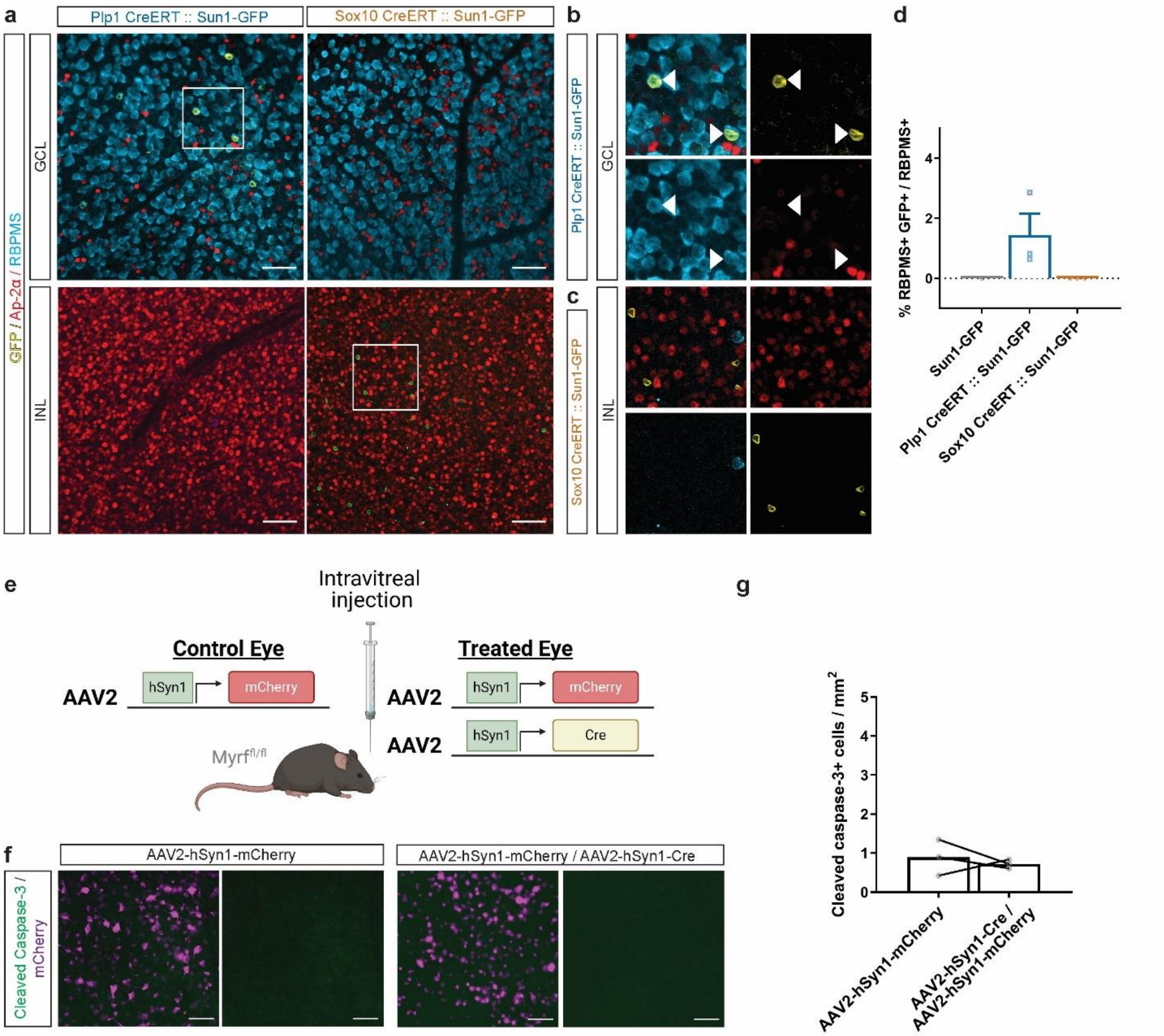
a
Plp1 CreERT :: Sun1-GFP
Sox10 CreERT :: Sun1-GFP
GFP / Ap-2α / RBPMS
GCL
INL
b
Plp1 CreERT :: Sun1-GFP
GCL
c
Sox10 CreERT :: Sun1-GFP
INL
d
% RBPMS+ GFP+ / RBPMS+
4
2
0
Sun1-GFP
Plp1 CreERT :: Sun1-GFP
Sox10 CreERT :: Sun1-GFP
e
Intravitreal injection
Control Eye
AAV2
hSyn1
mCherry
Treated Eye
AAV2
hSyn1
mCherry
AAV2
hSyn1
Cre
Myrf fl/fl
f
AAV2-hSyn1-mCherry
AAV2-hSyn1-mCherry / AAV2-hSyn1-Cre
Cleaved Caspase-3 / mCherry
g
Cleaved caspase-3+ cells / mm2
5
4
3
2
1
0
AAV2-hSyn1-mCherry
AAV2-hSyn1-Cre / AAV2-hSyn1-mCherry

**Supplementary Fig. 7: *Myrf* knockout from RGCs does not trigger apoptosis**.

**a** Images of the GCL and INL of the retina from Plp1 CreERT::Sun1-GFP and Sox10 CreERT::Sun1-GFP mice following tamoxifen administration. Boxed areas are shown **b** and **c**. **b** Plp1 CreERT::Sun1-GFP mice have occasional RBPMS+ cells that are GFP+ in the GCL. Arrowheads indicate co-labeled cells. **c** Sox10 CreERT::Sun1-GFP mice have recombined cells in the INL that do not express AP-2α, AP-2β or RBPMS. Sox10 CreERT::Sun1-GFP mice lack recombined cells in the GCL. **d** Quantification demonstrating only Plp1 CreERT:: Sun1 GFP mice have recombined RBPMS+ cells, which are absent in Sox10 CreERT:: Sun1 GFP mice. **e** Schematic of viral knockout approach using a AAV2-hSyn1-Cre expressing viruses to knock *Myrf* out of retinal neurons. **f** Retinae stained with cleaved caspase-3 and mCherry. Despite considerable viral infectivity evident by mCherry expression, there are very few apoptotic cells. **g** Quantification of apoptotic cells in the retinae of Myrf $^{fl/fl}$ following viral injection of AAV2-hSyn1-Cre/AAV2-hSyn1-mCherry or AAV2-hSyn1-mCherry alone. There is no statistical difference between treated and untreated eyes ($P$ = 0.6342). Paired t-test. Error bars are SEM. Scale bars are 50 µm in **a** and **f**. The schematic in **e** was created with BioRender.com.

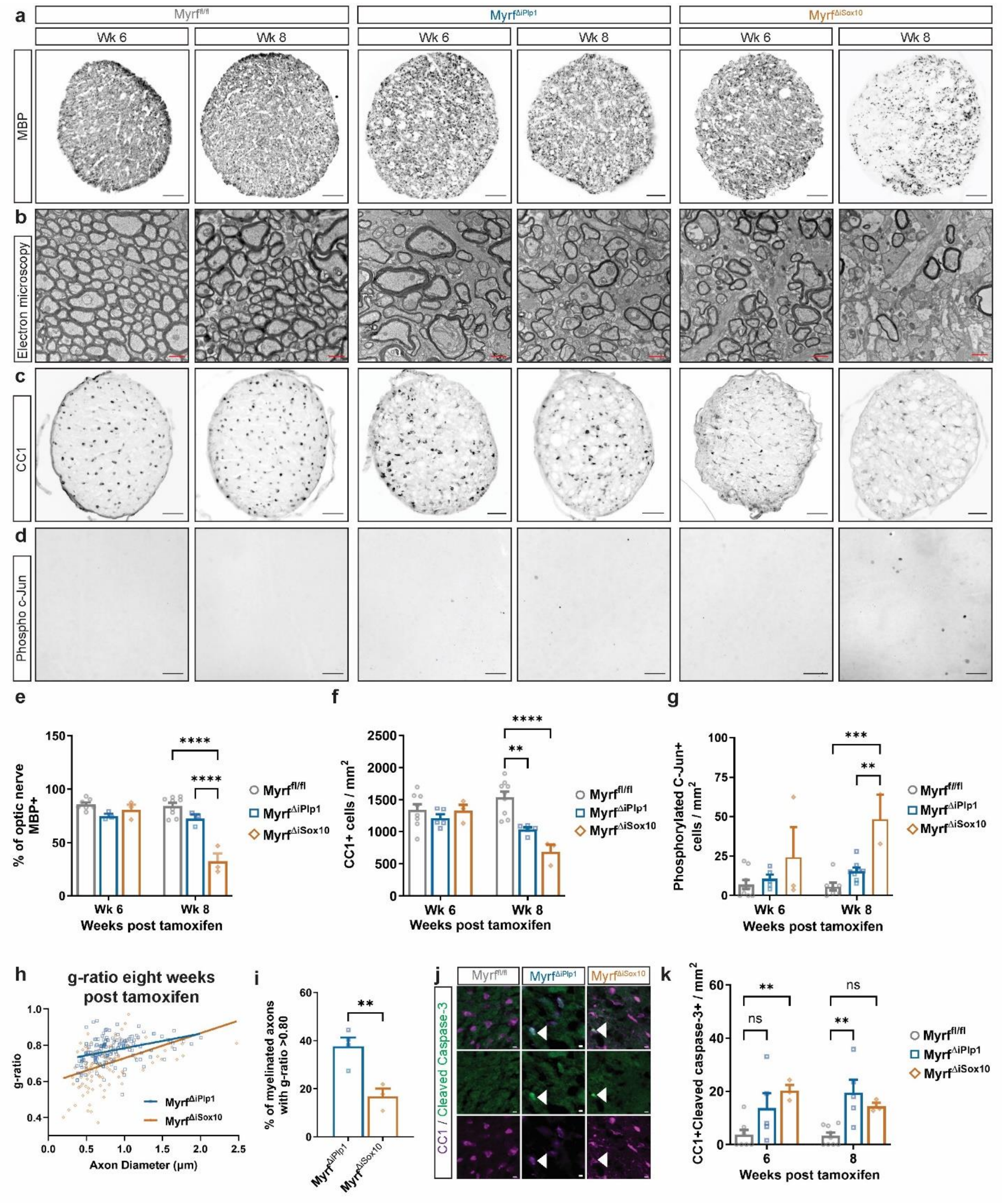
a
Myrf fl/fl
Myrf ΔiPlp1
Myrf ΔiSox10
Wk 6
Wk 8
MBP
b
Electron microscopy
c
CC1
d
Phospho c-Jun
e
% of optic nerve MBP+
Wk 6
Wk 8
Weeks post tamoxifen
****
Myrf fl/fl
Myrf ΔiPlp1
Myrf ΔiSox10
f
CC1+ cells / mm2
**
****
Weeks post tamoxifen
g
Phosphorylated C-Jun+ cells / mm2
***
**
Weeks post tamoxifen
h
g-ratio eight weeks post tamoxifen
g-ratio
Axon Diameter (µm)
Myrf ΔiPlp1
Myrf ΔiSox10
i
% of myelinated axons with g-ratio >0.80
**
Myrf ΔiPlp1
Myrf ΔiSox10
j
CC1 / Cleaved Caspase-3
k
CC1+Cleaved caspase-3+ / mm2
ns
**
Weeks post tamoxifen

**Supplementary Fig. 8: Demyelination and oligodendrocyte loss is coincident with phosphorylation of c-Jun in RGCs.**

**a** Overview images of MBP in the optic nerve of Myrf$^{fl/fl}$, Myrf$^{\Delta iPlp1}$ and Myrf$^{\Delta iSox10}$ mice at six and eight weeks post tamoxifen. **b** Electron micrographs reveal myelin delamination and degeneration at six weeks post tamoxifen and considerable demyelination by eight weeks post tamoxifen in Myrf$^{\Delta iSox10}$ mice. **c** Micrographs of optic nerve stained for the OL marker CC1. **d** Phosphorylated c-Jun in the retina of Myrf$^{fl/fl}$, Myrf$^{\Delta iPlp1}$ and Myrf$^{\Delta iSox10}$ mice. **e** Quantification of MBP within the optic nerve of Myrf$^{fl/fl}$, Myrf$^{\Delta iPlp1}$ and Myrf$^{\Delta iSox10}$ mice. There is a reduction of MBP by eight weeks post tamoxifen in Myrf$^{\Delta iSox10}$ mice relative to both Myrf$^{fl/fl}$ and Myrf$^{\Delta iPlp1}$ mice at ten weeks post tamoxifen ($p < 0.0001$). **f** Quantification of CC1+ OL density within the optic nerve at six and eight weeks post tamoxifen. There is a reduction in CC1+ cells in Myrf$^{\Delta iPlp1}$ ($P = 0.0011$) and Myrf$^{\Delta iSox10}$ mice ($P < 0.0001$) relative to Myrf$^{fl/fl}$ mice at eight weeks post tamoxifen. **g** Quantification of phosphorylated c-Jun in the retina of Myrf$^{fl/fl}$, Myrf$^{\Delta iPlp1}$ and Myrf$^{\Delta iSox10}$ mice at six and eight weeks post tamoxifen. There is an increase in the phosphorylation of c-Jun in Myrf$^{\Delta iSox10}$ relative to Myrf$^{\Delta iPlp1}$ ($P = 0.0045$) and Myrf$^{fl/fl}$ ($P = 0.0003$) at eight weeks post tamoxifen. **h** G-ratio at eight weeks post tamoxifen in Myrf$^{\Delta iPlp1}$ and Myrf$^{\Delta iSox10}$ mice. The average g-ratio is increased in Myrf$^{\Delta iPlp1}$ relative to Myrf$^{\Delta iSox10}$ mice. Student's t-test, $t = 3.358$, df = 7, $P = 0.0163$. $n$ = 3-4 mice per genotype, as shown. **i** Percentage of axons with a g-ratio > 0.8 in the optic nerve of Myrf$^{\Delta iPlp1}$ and Myrf$^{\Delta iSox10}$ mice at eight weeks post tamoxifen. These axons have thin myelin normally not found in the optic nerve. Myrf$^{\Delta iPlp1}$ mice have a higher percentage of thinly myelinated axons in their optic nerve relative to Myrf$^{\Delta iSox10}$ mice ($P = 0.0093$). **j** Images of cleaved caspase-3 along with CC1+ in Myrf$^{fl/fl}$, Myrf$^{\Delta iPlp1}$ and Myrf$^{\Delta iSox10}$ mice at six weeks post tamoxifen. Arrowheads indicate cleaved caspase-3+ CC1+ cells. **k** There is an increase in cleaved caspase-3+CC1+ OLs in Myrf$^{\Delta iPlp1}$ at eight weeks post tamoxifen ($P = 0.0013$) and in Myrf$^{\Delta iSox10}$ ($P = 0.0068$) at six post tamoxifen relative to Myrf$^{fl/fl}$. Myrf$^{\Delta iPlp1}$ and Myrf$^{\Delta iSox10}$ mice do not differ from each other at six ($P = 0.4535$) or at 8 ($P = 0.6135$) weeks post tamoxifen. Two-way ANOVA with Tukey's *post hoc* for individual comparisons in **e**, **f**, **g** and **k** and Student's t-test in **i**. Scale bars are 50 µm in **a**, **c**, **d**, 5µm in **j** and 1µm in **b**.

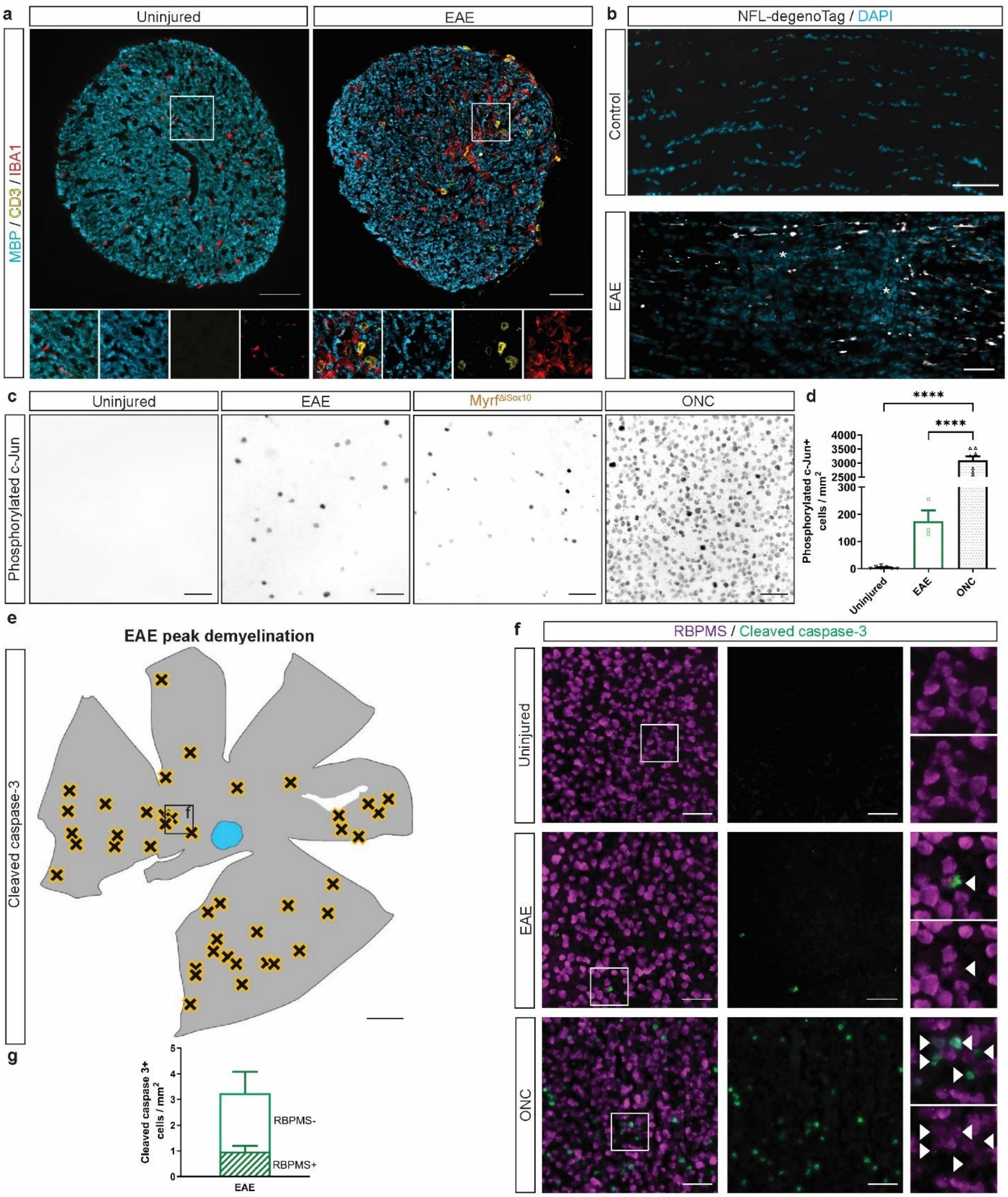

a
Uninjured
EAE
MBP / CD3 / IBA1
b
NFL-degenoTag / DAPI
Control
EAE
c
Uninjured
EAE
Myrf^ΔiSox10
ONC
Phosphorylated c-Jun
d
****
****
Phosphorylated c-Jun+ cells / mm²
4000
3500
3000
2500
300
200
100
0
Uninjured
EAE
ONC
e
EAE peak demyelination
Cleaved caspase-3
f
RBPMS / Cleaved caspase-3
Uninjured
EAE
ONC
g
Cleaved caspase 3+ cells / mm²
5
4
3
2
1
0
RBPMS-
RBPMS+
EAE

**Supplementary Fig. 9: Autoimmune-mediated demyelination triggers axonal damage, phosphorylation of c-Jun and apoptosis of RGCs.**

**a** Optic nerves from control and EAE mice stained with MBP for myelin, CD3 for T cells and IBA1 for microglia/macrophages. Boxed areas are enlarged below. **b** There is expression of an epitope of neurofilament light chain (NFL) known to be exposed during injury and that is found in EAE mice, which is often associated with higher densities of nuclei (likely inflammatory lesions, indicated by an asterisk). **c** Micrographs of the retina stained with phosphorylated c-Jun in uninjured mice, mice with EAE at peak disease, Myrf$^{\Delta iSox10}$ mice and mice three days post optic nerve crush (ONC). **d** Quantification of phosphorylated c-Jun in uninjured mice, EAE mice during peak disease and in mice with an ONC. ONC mice have increased phosphorylated c-Jun over both uninjured and EAE mice ($P < 0.0001$). EAE mice have a similar level of phosphorylation of c-Jun relative to Myrf$^{\Delta iSox10}$ (see Fig. 5g for comparison). Welch's ANOVA with Dunnett's T3 *post hoc* test. **e** Overview of a retina in a mouse with EAE during peak disease showing the location of cleaved caspase-3+ cells in the GCL. **f** High magnification micrographs of RBPMS and cleaved caspase-3+ cells. Arrowheads indicate cleaved caspase-3+ cells. Cleaved caspase-3+ cells often have weak RBPMS staining. **g** Density of cleaved-caspase-3+ cells ± RBPMS in EAE. Error bars are SEM. Scale bars are 50 µm in **a**- **c**, and **f** and 500 µm in **e**.

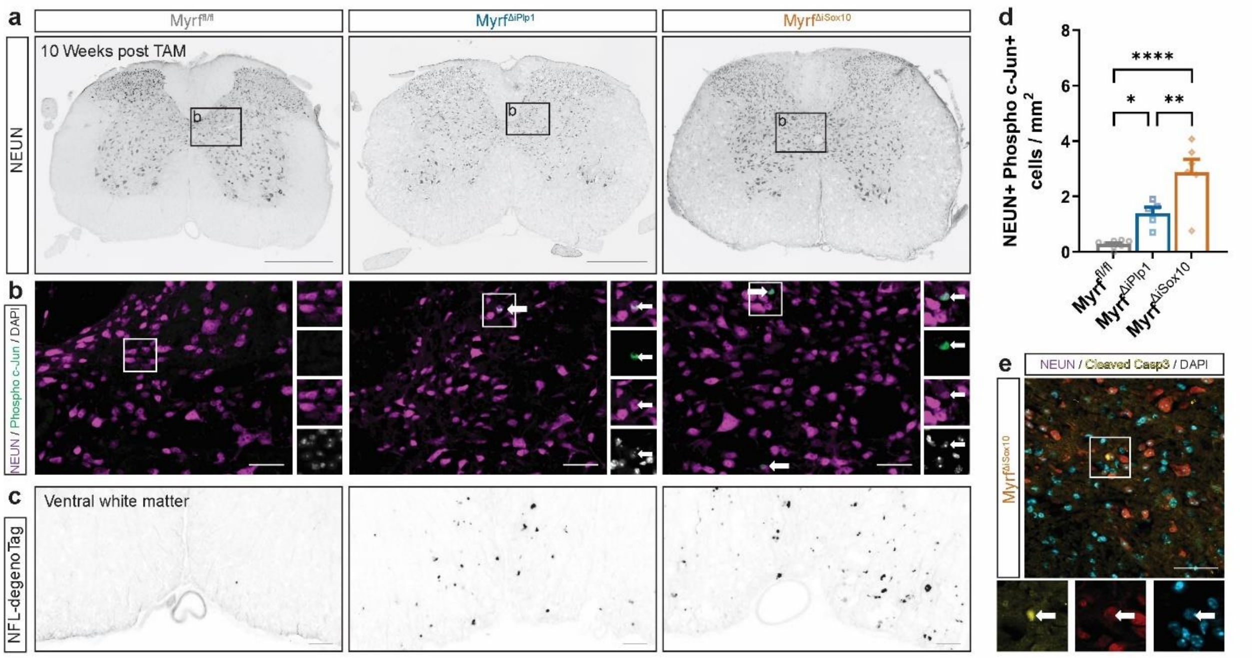


**Supplementary Fig. 10 Elevated phosphorylation of c-Jun in spinal neurons following demyelination in Myrf$^{\Delta iSox10}$ mice**

**a** Overview images of the lumbar spinal cord stained with NEUN. Boxed areas are shown in **b**. **b** Magnified images of the spinal cord grey matter stained with NEUN to label neurons and phosphorylated c-Jun. Boxed areas are shown to the right with individual channels. Arrows indicate NEUN+ phosphorylated c-Jun+ nuclei in Myrf$^{\Delta iSox10}$ and Myrf$^{\Delta iPlp1}$ mice. **c** Ventral white matter stained for NFL-degenoTag demonstrating damaged axons in both Myrf$^{\Delta iSox10}$ and Myrf$^{\Delta iPlp1}$ mice. **d** Myrf$^{\Delta iSox10}$ of mice show an increase in spinal neurons with phosphorylated c-Jun relative to both Myrf$^{\Delta iPlp1}$ ($P$ = 0.0068) and Myrf$^{fl/fl}$ ($P$ < 0.0001). Myrf$^{\Delta iPlp1}$ mice have increased density of phosphorylated c-Jun+ neurons ($P$ = 0.0300) relative to Myrf$^{fl/fl}$ mice. One-way ANOVA with Tukey's *post hoc* test for pairwise comparisons. **e** Example of a cleaved caspase-3+ neuron (NEUN+) in the spinal cord of Myrf$^{\Delta iSox10}$ mice. Error bars are SEM. Scale bars are 500 µm in **a** and 50 µm in **b, c and e**.

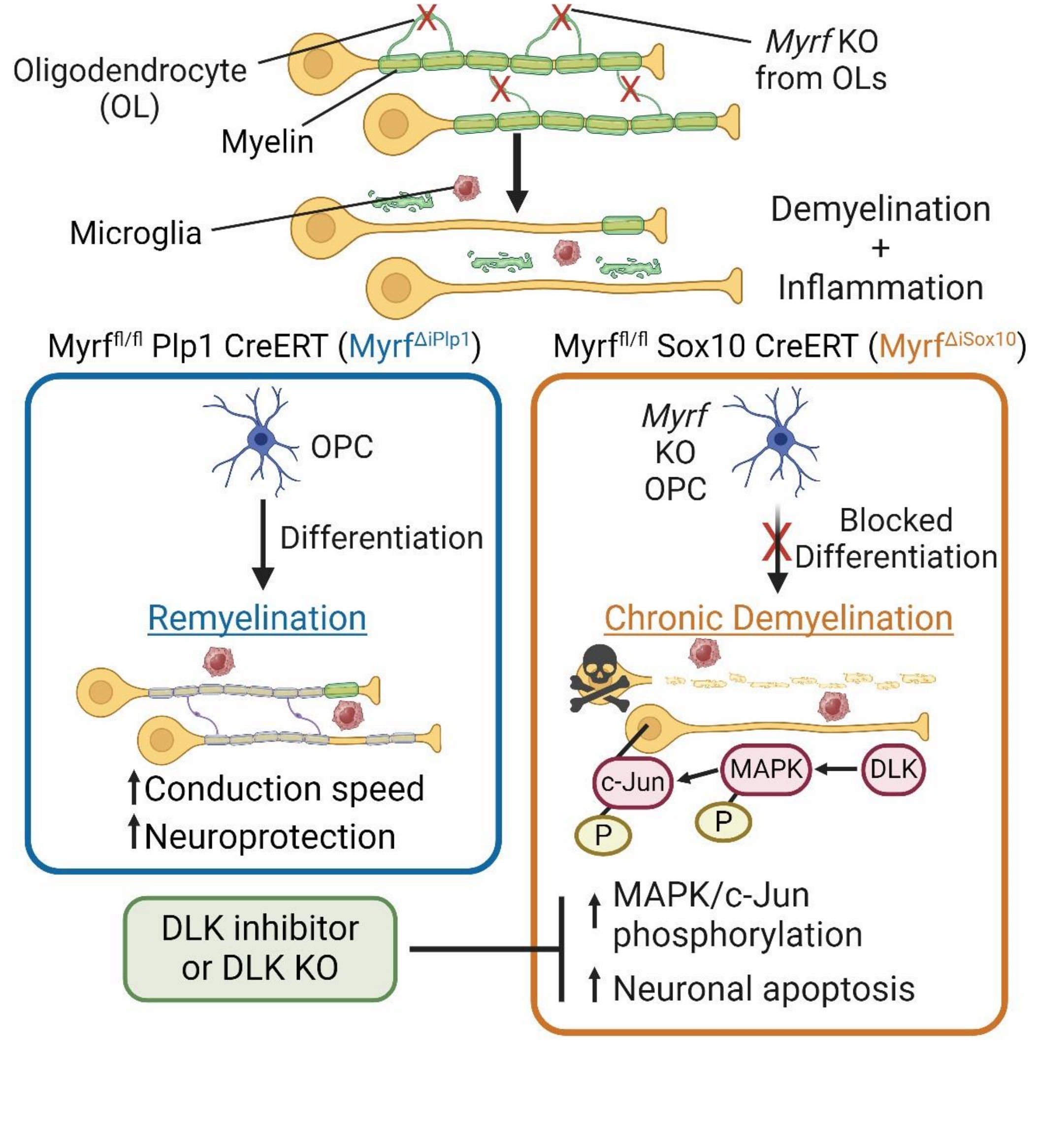


**Supplementary Fig. 11. DLK activation in chronically demyelinated drives neuronal apoptosis.**

Overview schematic with proposed model that a failure to remyelinate in Myrf $^{\Delta iSox10}$ mice is associated with DLK-mediated MAPK/c-Jun phosphorylation and apoptosis of the neuron. Application of DLK inhibitors or knockout of DLK are both able to block apoptosis in Myrf $^{\Delta iSox10}$ mice during chronic demyelination. Likewise, successful remyelination in Myrf $^{\Delta iPlp1}$ mice is associated with less activation of MAPK and c-Jun relative to mice without remyelination, and ultimately Myrf$^{\Delta iPlp1}$ do not feature neurodegeneration. The schematic was created with BioRender.com.

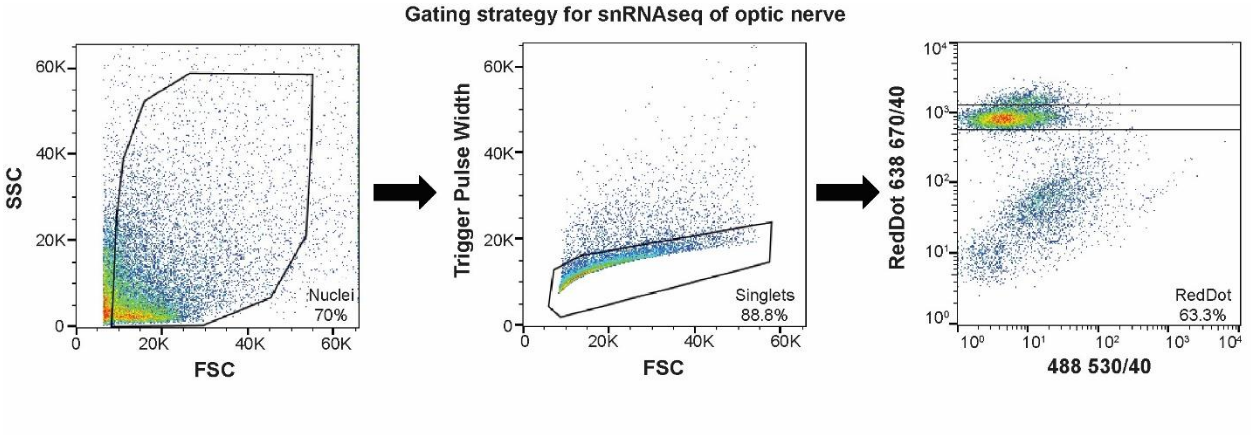


Supplementary Fig. 12 Gating strategy for nuclei for snRNAseq.

Nuclei were sorted for singlets and expression of RedDot nuclear dye. Example sort with gates shown.
